# Structural insights into assembly and function of the RSC chromatin remodeling complex

**DOI:** 10.1101/2020.03.24.006361

**Authors:** Richard W. Baker, Janice M. Reimer, Peter J. Carman, Tsutomu Arakawa, Roberto Dominguez, Andres E. Leschziner

**Affiliations:** Department of Cellular and Molecular Medicine, School of Medicine, University of California, San Diego, La Jolla, California 92093, USA; Section of Molecular Biology, Division of Biological Sciences, University of California San Diego, La Jolla, California 92093, USA; Department of Physiology, Perelman School of Medicine, University of Pennsylvania, Philadelphia, PA 19104-6085 USA; Alliance Protein Laboratories, a Division of KBI BioPharma, 6042 Cornerstone Court West, San Diego, CA 92121, USA; Department of Biochemistry and Biophysics, School of Medicine, University of North Carolina, Chapel Hill, Chapel Hill, NC, USA; Lineberger Comprehensive Cancer Center, University of North Carolina, Chapel Hill, NC, USA

## Abstract

Chromatin remodelers regulate the position and composition of nucleosomes throughout the genome, producing different remodeling outcomes despite a shared underlying mechanism based on a conserved RecA DNA translocase. How this functional diversity is achieved remains unknown despite recent cryo-electron microscopy (cryo-EM) reconstructions of several remodelers, including the yeast RSC complex. To address this, we have focused on a RSC subcomplex comprising its ATPase (Sth1), the essential actin-related proteins (ARPs) Arp7 and Arp9, and the fungal-specific protein Rtt102. Combining cryo-EM and biochemistry of this subcomplex, which exhibits regulation of remodeling by the ARPs, we show that ARP binding induces a helical conformation in the HSA domain of Sth1, which bridges the ATPase domain with the bulk of the complex. Surprisingly, the ARP module is rotated by 120° in the subcomplex relative to full RSC about a pivot point previously identified as a regulatory hub in Sth1, suggesting that large conformational changes are part of Sth1 regulation and RSC assembly. We also show that an interaction between Sth1 and the nucleosome acidic patch, which appears to be conserved among SWI/SNF remodelers, enhances remodeling. Taken together, our structural data shed light on the assembly and function of the RSC complex.

## Introduction

The position and composition of nucleosomes throughout the genome are tightly regulated. Key players in this regulation are the so-called chromatin remodelers, which non-covalently modify nucleosome architecture using a conserved ATPase subunit^1^. Chromatin remodeling is involved in nearly every aspect of transcriptional regulation and must therefore be controlled in a cell-cycle, cell-type, and developmentally specific manner^2^. To achieve this high degree of regulation, cells have evolved multiple families of remodelers, with each catalyzing distinct remodeling events, including sliding, ejection, and histone variant exchange. At the heart of each remodeler is an ATPase subunit with a conserved RecA domain, which couples ATP hydrolysis to translocation along a single strand of DNA. Domains flanking the core ATPase as well as accessory subunits allow each family to use the same biochemical RecA function to produce diverse remodelling outcomes. It is not yet fully understood how this functional diversity is achieved.

The SWI/SNF family of chromatin remodelers, which in yeast includes the eponymous SWI/SNF complex and the RSC complex, modify the position and spacing of nucleosomes at promoter regions throughout the genome and are capable of ejecting histone octamers from the DNA^3,4^. In yeast, RSC is a 17-subunit complex with a MW of > 1 MDa^5,6^, and its subunit architecture and biological function are broadly conserved from yeast to humans. Recent meta-genomic analysis has revealed that SWI/SNF family genes in humans are highly mutated in cancer^7–9^, with mutation rates of all genes in aggregate (19%) rivaling those of *TP53* (26%)^9,10^. Most of the mutations lie in the accessory subunits of the complex, yet oncogenesis has in at least one case been shown to require an active ATPase subunit^11^. This suggests that not only are accessory subunits fundamental components of the remodeling process itself, but that misregulation of remodeling is a hallmark of human disease.

Significant genetic and biochemical insight has been gained by focusing on the essential actin-related proteins (ARPs) Arp7 and Arp9, which are conserved in all SWI/SNF remodelers from yeast to humans^12,13^. Deletion of the *ARP7* or *ARP9* genes is lethal in yeast ^14,15^. The “ARP module”, which consists of Arp7, Arp9, and the fungal-specific protein Rtt102, binds to the HSA domain in Sth1, a helical segment N-terminal to the ATPase domain^15–17^. Binding of the ARP module lowers the intrinsic rate of ATPase turnover of Sth1, yet paradoxically increases remodeling rates, suggesting that ARP binding acts to convert ATP turnover by the core ATPase into productive remodelling events^18^. Additionally, two other SWI/SNF-specific domains, the post-HSA and Protrusion (P1), seem to interact both physically^17,19^ and genetically^15^ with the HSA domain, indicating that they act as a regulatory hub to control ATPase turnover and remodelling^18^. Removal of the ARPs from the full RSC complex results in basal remodelling roughly equivalent to that of Sth1 alone, suggesting that they perform a core function in activating remodeling by RSC^18^. A mechanistic understanding of these regulatory interactions is still missing as the recent cryo-EM structures of the yeast RSC^20–22^ and SWI/SNF^23^ complexes, and the BAF complex, the human ortholog of SWI/SNF ^24^ did not reach high enough resolution in the ARP module or the HSA/post-HSA/P1 regulatory hub. To bridge this gap, we focused on the well-characterized Sth1/Arp7/Arp9/Rtt102 RSC sub-complex (Fig.1a) that has been used to gain much of our current understanding of the regulation of remodeling in RSC ^17,18,25–27^.

**Fig. 1.**
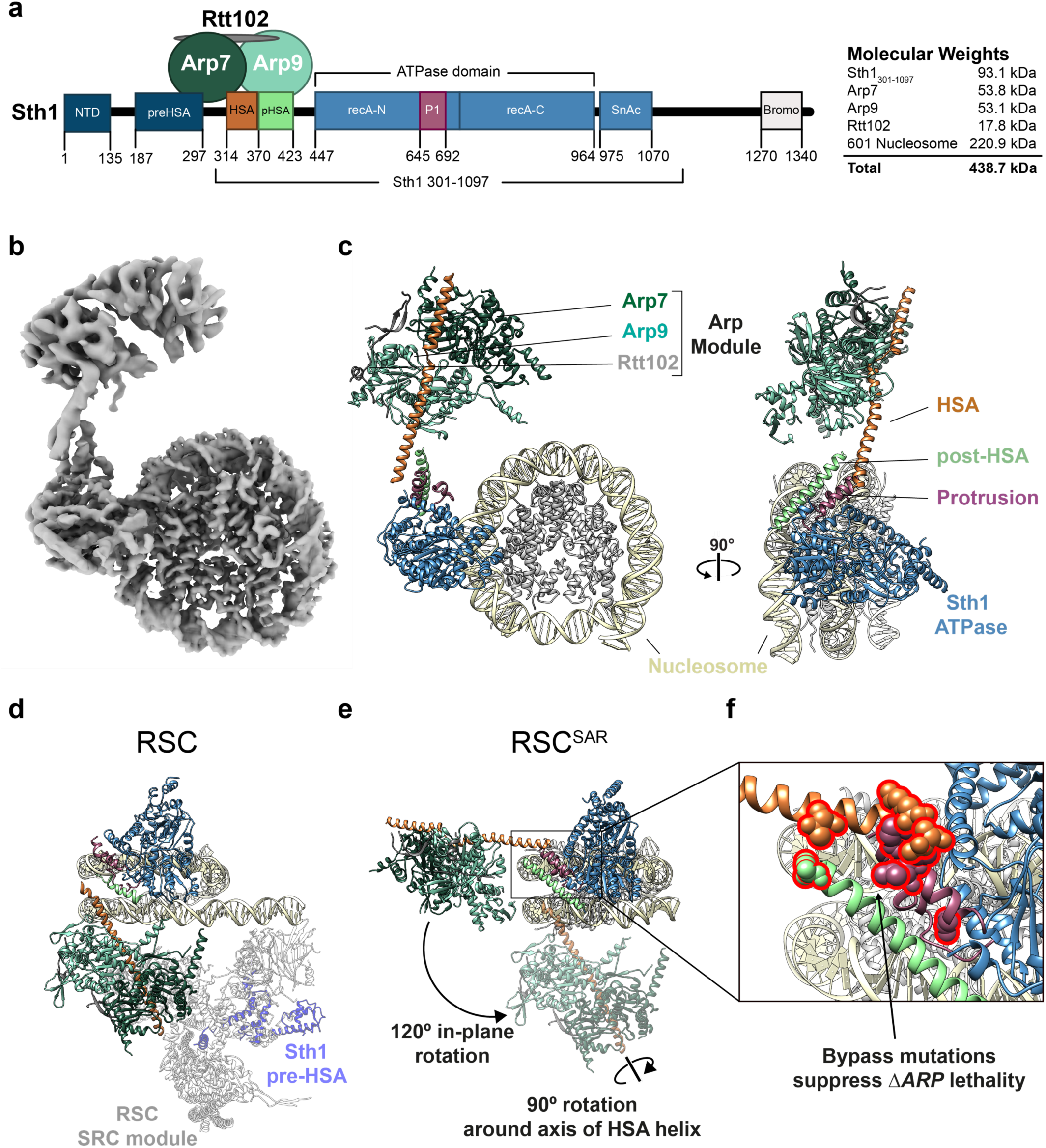
Cryo-EM structure of the Sth1-Arp7-Arp9-Rtt102 RSC subcomplex (RSC^SAR^) bound to a nucleosome. **a**, Schematic representation of RSC^SAR^, with Sth1 domains and their boundaries indicated. The same color scheme is used in all figures. **b**, 3.9 Å Cryo-EM map of RSC^SAR^ bound to a nucleosome. **c**, Molecular model of the RSC^SAR^:nucleosome complex. **d, e**, The position of the ARP module differs by 120° between RSC^SAR^ and RSC. **d**, A recently published structure of a RSC:nucleosome complex (6KW3.pdb^22^) with the portion corresponding to RSC^SAR^ shown with the same colors introduced above. The N-terminal portion of Sth1 preceding the HSA, which is absent in the Sth1 construct used in RSC^SAR^, is shown in dim purple. The Substrate Recognition Complex (SRC) of RSC is shown in grey. **e**, RSC^SAR^ is shown with the nucleosome and Sth1 in the same orientation as that in (d). The position occupied by the ARP module in RSC (d) is shown in dim colors. The movements required to convert the ARP module from its position in RSC^SAR^ to that in RSC—a 120° in-plane rotation and a 90° rotation about the HSA helix—are indicated. **f**, Close up of the regulatory hub located at the base of the HSA helix, consisting of the C-terminal end of the HSA helix, the post-HSA and P1. Residues where bypass mutations rescue *ΔARP* lethality are shown in space filling representation.

We present here a cryo-EM reconstruction of the <u>S</u>th1/<u>A</u>rp7/<u>A</u>rp9/<u>R</u>tt102 complex (which we will refer to as “RSC^SAR^”) bound to a nucleosome in the presence of the ATP analog ADP beryllium fluoride (ADP-BeF_3_). Using circular dichroism (CD) spectroscopy, we show that binding of the ARP module induces the helical folding of the HSA domain in Sth1. Our structure revealed that the ordered HSA domain physically interacts with the post-HSA and P1 domains, forming a pseudo-four helix bundle. Importantly, mutations that bypass *ΔARP* lethality in yeast cluster to this helical bundle, suggesting that ARP binding is transmitted to the Sth1 enzyme through this regulatory hub by inducing the folding of the HSA helix. Surprisingly, we observed a 120° rotation of the ARP module in RSC^SAR^ compared to the full RSC complex, with the regulatory hub serving as its hinge point. We propose that binding of the ARP subunits creates a rigid “rod” that communicates conformational changes between the HSA/post-HSA/P1 regulatory hub of the Sth1 ATPase and the bulk of the RSC complex. Our cryo-EM analysis also identified a conserved domain in SWI/SNF family enzymes that interacts directly with the nucleosome acidic patch and helix 1 on H2B, and functional assays showed that disrupting this interaction significantly reduced remodeling. Finally, our structure showed that the RSC^SAR^ complex can peel away DNA at the nucleosomal exit site, a function previously attributed to the full complex.

## Results

### Cryo-EM structure of a regulated RSC subcomplex

Recent cryo-EM reconstructions of the RSC^20–22^, SWI/SNF^23^ and BAF^24^ complexes revealed, for the first time, the overall architecture of these important remodelers. However, these structures did not provide a mechanistic basis for how accessory subunits, in particular the ARPs, regulate the core ATPase domain in Sth1 due to the limited resolutions in that part of the maps. The BAF-nucleosome structure, which shows the strongest ARP density among the different structures, has a resolution of 9-15 Å in this region; focused refinement was used in that work to improve the density of the ARPs and the HSA helix but this analysis excluded the interface of the HSA domain with the ATPase. To address how accessory subunits regulate the catalytic ATPase domain, we turned to a well-studied RSC sub-complex that can remodel nucleosomes and shows regulation of its remodeling activity by the ARPs^17,18,25,26,28^. This complex includes a truncated form of Sth1 (AA 301-1097) and three subunits—Arp7, Arp9 and Rtt102—that constitute the “ARP module”, accounting for ∼25% of the total mass of RSC (Figure 1a). We reasoned that this simpler system might allow us to obtain higher resolutions at the ATPase/ARP interface and thus shed more light on the regulation of Sth1 by the ARPs. We refer to this complex as RSC^SAR^ throughout the manuscript (SAR = <u>S</u>th1 + <u>A</u>RPs + <u>R</u>tt102).

To understand the structural consequences of ARP-module binding, we sought to determine cryo-EM structures of RSC^SAR^ both alone and bound to the nucleosome. We obtained a 4.2 Å structure of RSC^SAR^ by itself, showing well-ordered density for the ARP module (Sth1_HSA_/Arp7/Arp9/Rtt102), but lacking density for the ATPase domain of Sth1. We used our map to build a model of Sth1_HSA_/Arp7/Arp9/Rtt102 (Supp. Fig 1) which shows differences in the HSA domain when compared to the Snf2_HSA_/Arp7/Arp9/Rtt102 crystal structure^16^ (Supp. Fig. 1d). In the crystal structure, the Snf2 HSA domain binds to Arp7/Arp9 as a single α-helix, whereas the Sth1 HSA in our cryo-EM structures forms two helices connected by a short loop. Whether this structural difference between SWI/SNF and RSC has any functional significance is unclear.

We next determined a cryo-EM structure of RSC^SAR^ bound to the nucleosome in the presence of Mg^2+^ and ADP-BeF_3_ at a nominal resolution of 3.9 Å (Fig. 1b). Initial reconstructions showed strong density for the Sth1 ATPase domain and the nucleosome, and clear, albeit fragmented, density near the N-terminus of Sth1 (Supp. Fig 2). Using 3D classification, followed by multi-body refinement, we were able to improve the resolution of the map (Nucleosome: 3.7 Å; ATPase domain: 4.0 Å; ARP module: 6.5 Å) and unambiguously assign all components of the RSC^SAR^ complex (Fig 1c, Supp. Fig 2-4). We built a model of RSC^SAR^-nucleosome using the cryo-EM structure of Snf2-nucleosome and our Sth1_HSA_/Arp7/Arp9/Rtt102 model (Supp. Fig 4a-d). Overall, our RSC^SAR^-nucleosome structure shows a RecA ATPase bound to the nucleosome at superhelical location 2 (SHL2), consistent with previous cryo-EM structures^19,29,30^. Surprisingly, the ARP module extends from the N-terminus of the ATPase domain, lying roughly on the same plane as the nucleosome (Fig. 1b,c). This is in contrast to the structure of the full RSC complex, as well as those of the SWI/SNF and BAF complexes, where the HSA and associated ARP module are rotated by ∼120° relative to their position in RSC^SAR^ (Fig. 1d,e and Supp. Fig. 5)^20–24^. The C-terminal end of the HSA helix contacts the ATPase and engages with the post-HSA and P1 domains, as well as the Brace Helices (Fig. 1e and Fig. 2). The importance of these domains, which act as a regulatory hub within the ATPase domain, is discussed below.

**Fig. 2.**
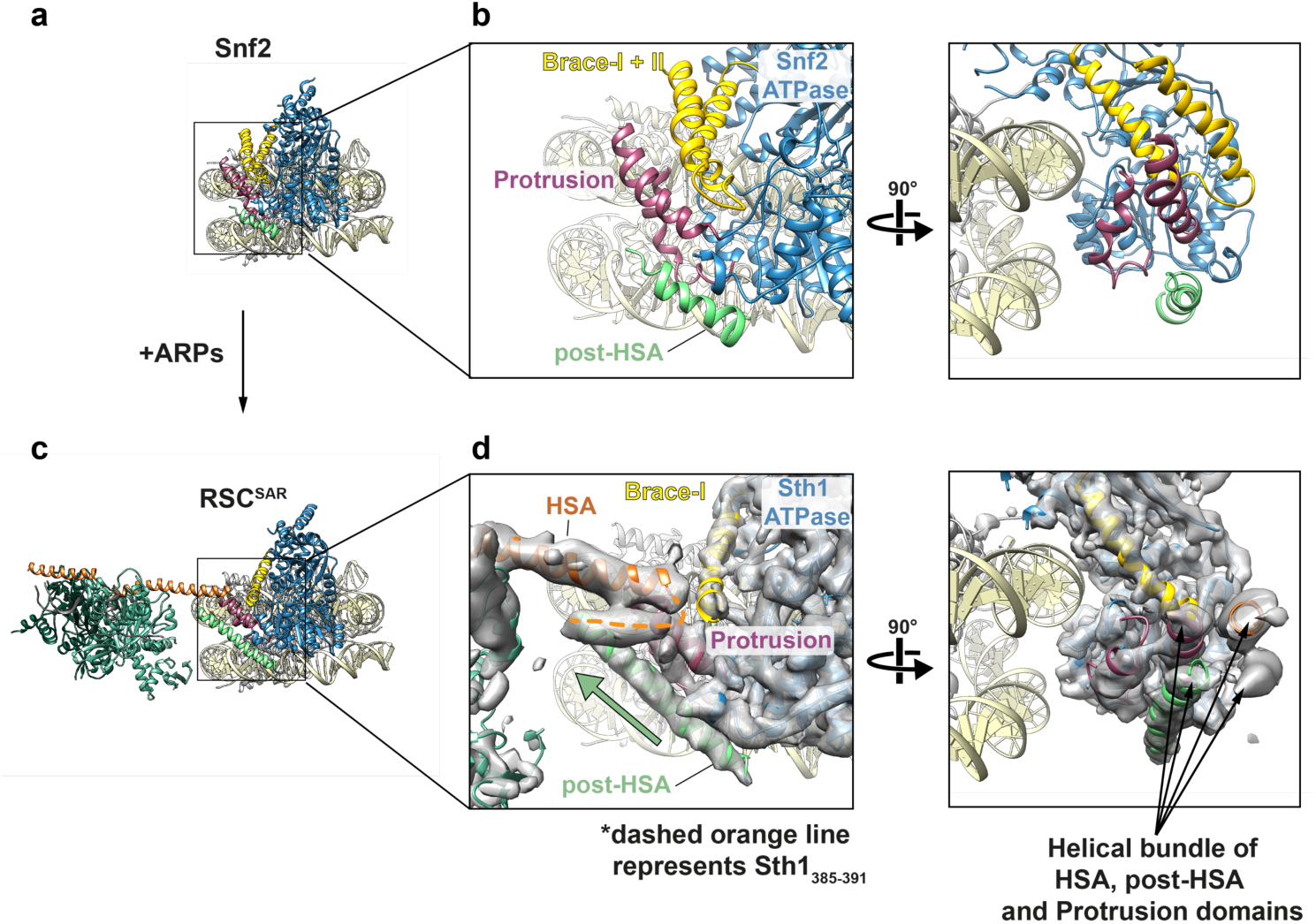
ARP binding produces structural changes in the HSA/post-HSA/P1 regulatory hub. **a**, Previously published structure of Snf2 bound to the nucleosome in the presence of ADP-BeF_3_^29^. Tan: nucleosome; Blue: ATPase domain; Yellow: Brace Helices; Dark pink: P1 domain; Green: post-HSA domain. **b**, Two views showing the interaction of post-HSA/P1 domains and the Brace Helices. **c**, Structure of RSC^SAR^ bound to the nucleosome in the presence of ADP-BeF_3_ (this work). Coloring is the same as in panel (a) plus Orange: HSA domain; Dark Green: Arp7; Light Green: Arp9; Grey: Rtt102. **d**, Equivalent views as those in (b), showing the interaction of HSA/post-HSA/P1 domains and the Brace Helices. The cryo-EM density for RSC^SAR^ is included in this panel. A pseudo four-helix bundle is formed by the HSA/post-HSA/P1 domains. The dashed orange line indicates density connecting the HSA to the post-HSA. The green arrow highlights the longer post-HSA helix present in RSC^SAR^ vs. Snf2 (compare (b) and (d)).

### Assembly of the full RSC-nucleosome complex requires large conformational changes in the ARP module

In each of the recently reported structures of SWI/SNF remodelers (RSC, SWI/SNF, and BAF) bound to the nucleosome^20–24^ the ATPase is bound at SHL2 on the nucleosome while the rest of the complex resides on the distal side of the nucleosome (Fig. 1d and Supp. Fig. 5). The portion of RSC not included in our complex has been referred to as the Substrate Recognition Complex (SRC) and contains several DNA- and histone-binding domains that likely aid in determining RSC localization throughout the genome. The main connection between Sth1 and the SRC is the ARP module, which seems to have little interaction with the rest of the complex (Supp. Fig. 5). Notably, the ARP module itself occupies very different positions in RSC^SAR^ and the full RSC complexes: going from its orientation in RSC^SAR^ to that in full RSC involves a 120° rotation away from the plane of the nucleosome and a 90° rotation about the axis of the HSA helix (Fig. 1d,e). This results in a ∼80 Å displacement of the center of mass of the module between the two structures. When engaged in the full RSC complex, the ARP module extends perpendicularly from the plane of the nucleosome and serves as a rigid connection between the ATPase (Sth1) and the SRC module, which together engage both faces of the nucleosome (Fig. 1d and Supp. Fig. 5b).

Although we were able to build a full model of the RSC^SAR^ complex, we did observe significant conformational heterogeneity in our dataset. Our biggest improvement in resolution came from using multi-body refinement^31^ (Supp. Fig. 3), which refines user-defined portions of the model as independent bodies. This analysis showed that the ARP module undergoes conformational changes of upwards of ∼70 Å in RSC^SAR^, with the structures representing this motion forming a cone of conformations anchored at the N-terminus of the ATPase domain of Sth1, where the HSA domain engages two structural elements of the regulatory hub: the post-HSA and P1 domains (Supp. Fig. 3d). Importantly, focused 3D classification did not identify particles where the ARP module occupied the same position as that seen in the full RSC complex. While we cannot rule out the possibility that a small subset of RSC^SAR^ particles representing that conformation do exist in our dataset, our data suggest that the conformation of the ARP module in the RSC^SAR^ complex is distinct from that seen in full RSC. Thus, formation of the full RSC complex likely requires a large conformational change in the ARP module during assembly, upon engagement with the nucleosome, or both.

### The HSA helix interacts with a regulatory hub in the ATPase domain of Sth1

Previous data have shown that three accessory domains are essential to the function of Sth1–the HSA, post-HSA, and P1 domains^15^. To understand the consequences of ARP binding and RSC assembly on this important regulatory region, we compared cryo-EM structures of Snf2^29^, RSC^SAR^, and RSC^22^ (Fig. 2 and Supp. Figs. 4 and 6). The first cryo-EM structures of Snf2 bound to the nucleosome revealed that the post-HSA and P1 domains physically interact with each other (Fig. 2b). Our cryo-EM structure of RSC^SAR^ bound to a nucleosome showed that in the presence of the ARP module, the HSA helix interacts with the post-HSA and P1 domains, creating a pseudo four-helix bundle (Fig. 2d, right). Interestingly, mutations that bypass the need for the *ARP9* gene in yeast all cluster in this region^15^, and 8 out of 9 point mutants can be mapped onto ordered regions in RSC^SAR^ (Fig. 1f). Additionally, a truncation of AA 385-392 in Sth1 that was also found to suppress loss of *ARP9* corresponds to a region of the four-helix bundle linking the HSA and post-HSA domains (Fig. 2d). Compared with Snf2, we see an extension of the post-HSA helix by 5 helical turns (Fig2b,d and Supp. Fig. 6a) and a displacement of the P1 domain by ∼ 7 Å (Supp. Fig. 6b).

In addition to the HSA/post-HSA/P1 regulatory hub, SWI/SNF remodelers also contain a set of helices that link the two lobes of the ATPase, termed Brace Helix I and II^30^. The Brace Helices interact with the P1 domain and are also in close proximity to the ordered HSA domain in the RSC^SAR^ structure (Fig. 2d). While Brace Helix I shows strong density in our cryo-EM map, Brace Helix II is completely disordered (Supp. Fig. 4b,c). This conformational change is likely intimately associated with the nucleotide state of the ATPase, as both of these helices are ordered in the Snf2-apo and Snf2-ADP cryo-EM structures^29^ (Supp. Fig. 4e). Although Brace Helix II was modeled in the Sfn2-ADP-BeF_3_ structure (Supp. Fig. 4b,e)^29^, there is no ordered density in the corresponding part of the cryo-EM map (Supp. Fig. 4e). Because the Brace Helices physically link the two lobes of the ATPase, interaction of this motif with the HSA/post-HSA/P1 hub likely serves as a conduit to relay the conformation of the ATPase lobes to the rest of the RSC complex, and vice versa.

### Binding of the ARP module induces folding of the HSA and post-HSA domains

The HSA and post-HSA domains in RSC^SAR^ showed extended helical structures compared with those seen in the cryo-EM structures of the related Snf2 enzyme^19,29^ (Fig. 2b,d and Supp. Fig. 4e). Given that a major difference between our structure of RSC^SAR^ and those of the Snf2-nucleosome complex is the presence of the ARPs, and that the ARPs have no binding partners outside the HSA domain in either RSC or RSC^SAR^, we hypothesized that a key function of the ARPs might be to induce folding of the HSA domain into an alpha helix.

To determine if the HSA domain could adopt a helical conformation on its own, we performed Circular Dichroism (CD) spectroscopy on a peptide including the region of the HSA domain (AA 316-377) that was ordered in our cryo-EM reconstructions and in previous X-ray structures of the ARP module^16,17^. Our analysis showed that the HSA exhibits a random coil conformation in the absence of the ARP module (Fig. 3a, blue line). In contrast, comparing spectra of the ARP module with and without the HSA peptide (Fig. 3b) showed that addition of the ARP module induced a helical fold in the HSA domain (Fig. 3a, orange line).

**Fig. 3.**
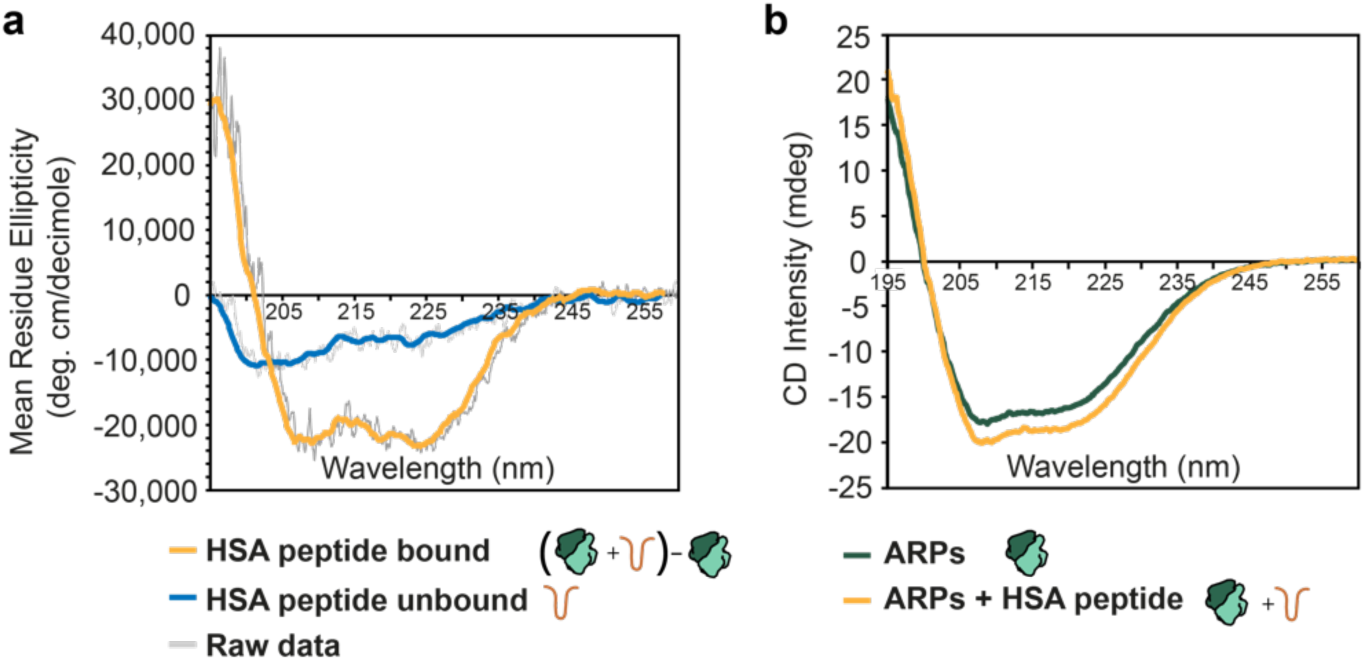
Binding of the ARP module induces folding of the HSA helix. **a**, Molar ellipticity plot for the HSA peptide bound to the ARP module (orange) or without ARP module (blue). Raw data are shown in grey, with the colored line representing a smoothed trace. Bound HSA data was calculated by subtracting the plots shown in (b) and calculating molar ellipticity based on the HSA peptide (see Methods). **b**, Circular dichroism plot for the ARP module (green) and the ARP module plus HSA peptide (orange). Raw, unsmoothed data are shown. All spectra were collected with identical protein concentrations and with complexes at equimolar ratios.

### RSC^SAR^ can break DNA:histone contacts at the exit site

The mechanism of step-wise DNA translocation by SNF2-family ATPase enzymes has recently been elucidated with cryo-EM structures of Snf2 bound to the nucleosome^29^. To understand how RSC^SAR^ (4 subunits) might distort the nucleosomal DNA relative to Snf2 (1 subunit), we compared our structure with that of Snf2-nucleosome in the ADP-BeF_3_ bound state (Fig 4, Supp. Fig. 6 and 7). We observed a near identical conformation of the nucleosomal DNA at SHL2 and at other super-helical sites throughout the nucleosomes (Supp. Fig. 7). The conformation of the ATPase domain of Sth1 in our RSC^SAR^ model is in good agreement with that of Snf2 (Supp. Fig. 6), except for the changes in the HSA, post-HSA, and P1 domains discussed above. However, we do see a large difference in the conformation of the DNA at the nucleosome exit site compared to that seen with Snf2. While all reported Snf2 cryo-EM datasets show a canonical nucleosomal DNA^19,29^, 26.5% of particles in our dataset have an exit site DNA peeled by approximately 40°, as was recently reported in structures of both RSC and SWI/SNF bound to the nucleosome ^21,22^ (Fig. 4 and Supp. Fig. 1). Our data suggest that peeling of the exit site DNA and breaking of multiple DNA-histone contacts is a feature of the RSC^SAR^ complex and does not require a full complement of RSC subunits, as has been proposed^22^. Additionally, previous biochemical analysis has shown that mutation of several basic residues in the N-lobe of Snf2, which lies in close proximity to the exit site DNA, reduces the remodeling rate of the isolated Snf2 enzyme^19^. Thus, peeling of the exit site DNA is likely an important component of the remodeling reaction and is mediated, at least in part, by the ATPase itself.

**Fig. 4.**
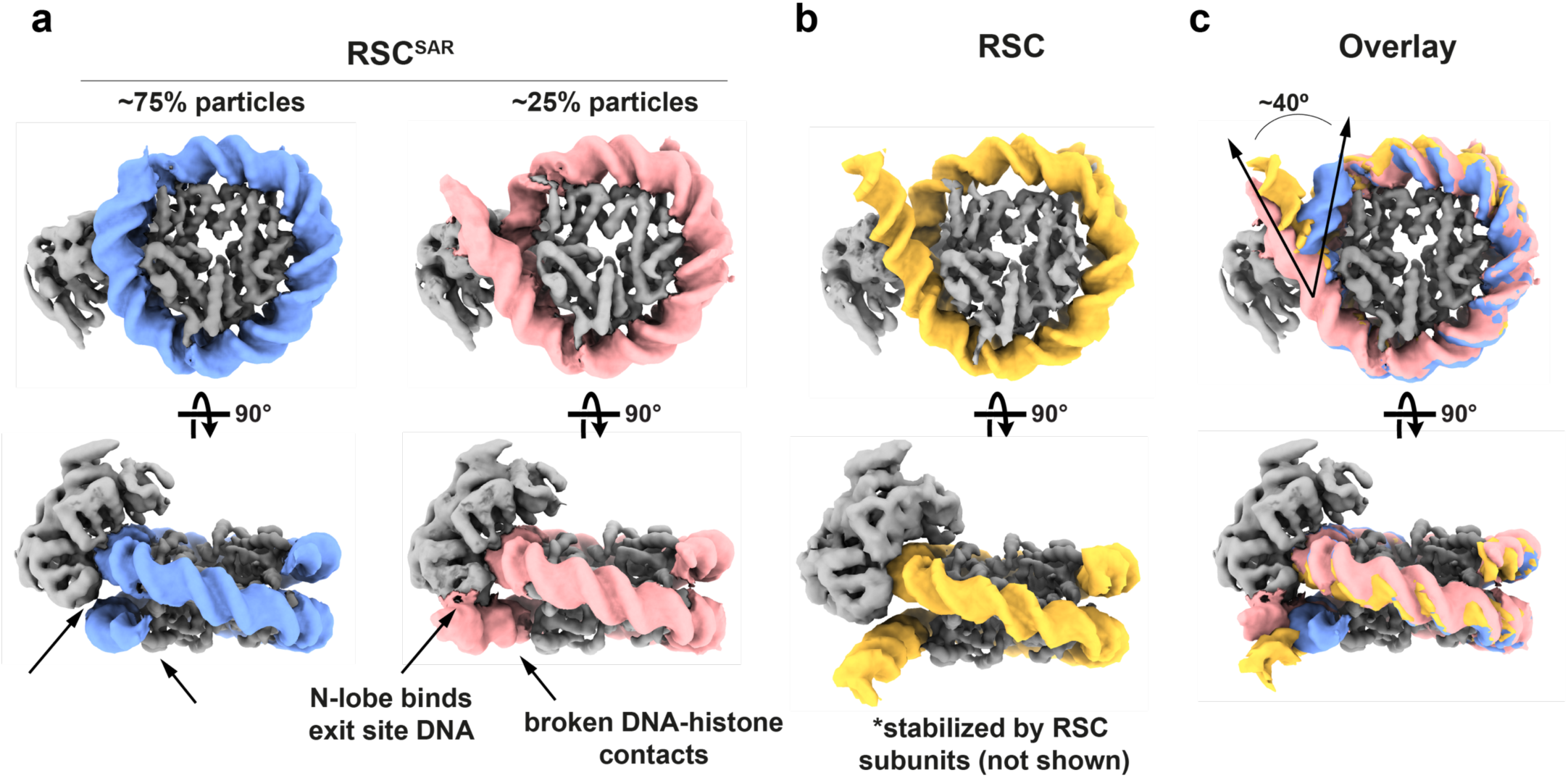
RSC^SAR^ peels off nucleosomal DNA at the exit site. **a**, 3D classification of RSC^SAR^ particles showed that ∼25% have a peeled DNA conformation at the DNA exit site. Two cryo-EM maps are shown representing these populations. Blue: nucleosomal DNA with unpeeled exit site, Pink: nucleosomal DNA with peeled exit site. **b**, The exit site DNA in the published RSC-nucleosome cryo-EM structure showed a peeled conformation that was stabilized by accessory subunits in RSC not present in RSC^SAR^. This panel shows a density generated from a PDB model; only the ATPase domain of Sth1 and the nucleosome were included to simplify the comparison with the structures in (a). Yellow: DNA from RSC-nucleosome model. **c**, An overlay of the three structures (RSC^SAR^ peeled; RSC^SAR^ unpeeled, and RSC) shows that RSC^SAR^ is able to induce a DNA conformation at the exit site equivalent to that of full RSC in the absence of the accessory subunits present in the latter. Colors equivalent to panels a and b.

### Sth1 uses a conserved domain to bind the nucleosome’s acidic patch and promote remodeling

During our initial cryo-EM processing we noticed that a C-terminal region of Sth1 extends from the ATPase domain and appears to make contact with the surface of the histone core (Fig. 5a, red dashed circle, and Supp. Fig. 8a,b). Due to the lower resolution of this part of the map, this density is more visible in maps that are not sharpened or are filtered by local resolution (Supp. Fig. 8a,b). This Sth1 density seems to directly interact with a region of the nucleosome known as the acidic patch, in agreement with a similar density observed in cryo-EM reconstructions of the full RSC complex^21^. The nucleosome acidic patch is composed of 8 acidic residues found on histones H2A and H2B and is used by many proteins to interact with the nucleosome^32–35^ (Fig. 5b). The characteristic binding motif for the acidic patch is an arginine anchor that makes hydrogen bonds with H2A residues E61, D90, and D92 (Fig. 5c). We observed a pocket of unassigned density directly above the acidic patch that corresponds with the binding site for the arginine anchor (Fig. 5a, red circle and 5c). Sth1 contains an arginine-rich area (which we will refer to as the “basic patch”) following the SnAc domain (Fig. 6a), and the distance from the last modeled residue of the SnAc domain would place the basic patch in reasonable proximity to the acidic patch (31 residues to span ∼40 Å distance). Furthermore, this C-terminal basic patch (Sth1 residues 1084-1096) is conserved in SWI/SNF remodelers, but not in other remodeler families (Fig. 6a).

**Fig. 5.**
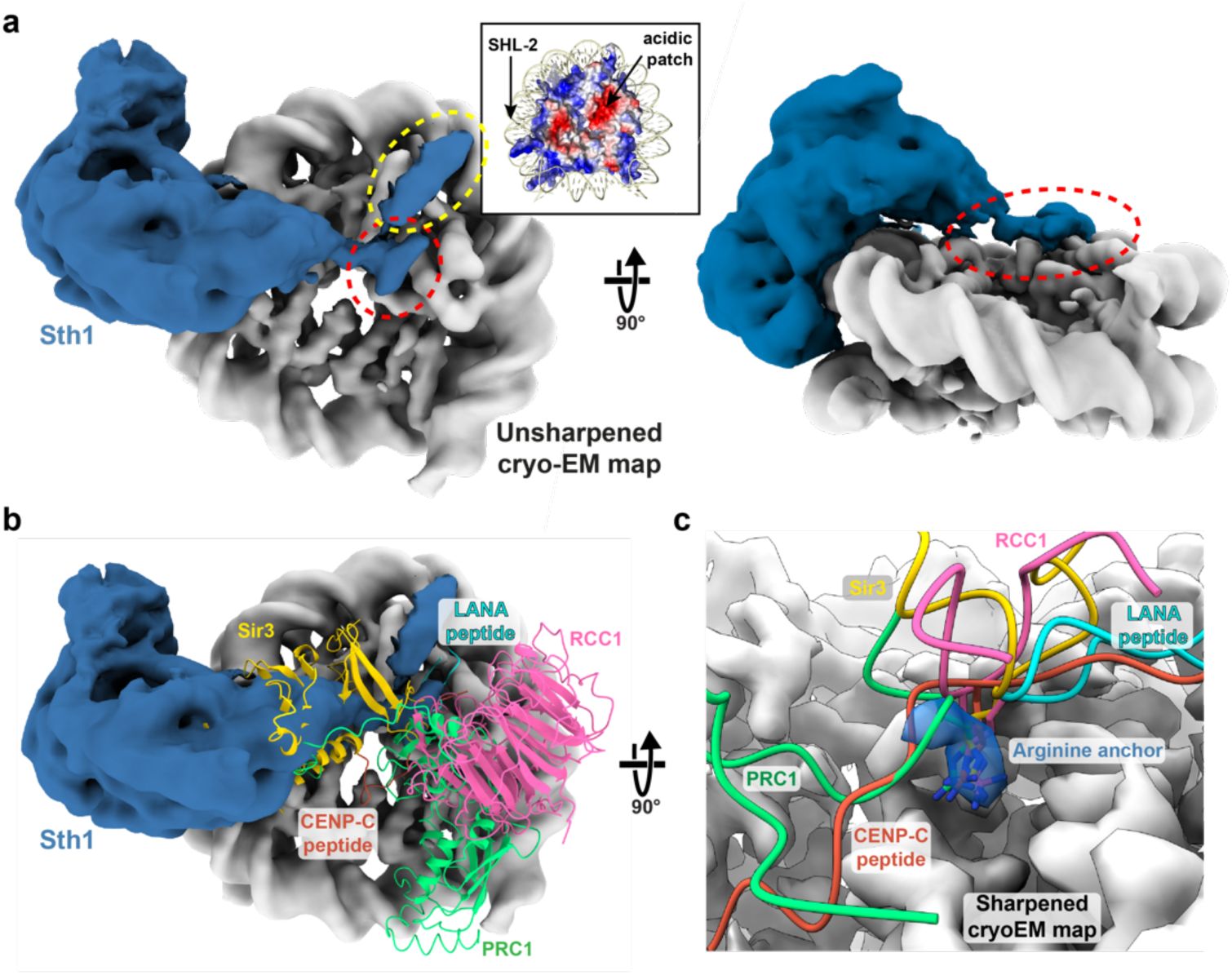
A basic stretch in Sth1 recognizes the nucleosome acidic patch. **a**, Unsharpened cryo-EM map of RSC^SAR^. The map was manually segmented and colored grey for the nucleosome and blue for Sth1. Inset: the nucleosome is shown colored by electrostatic potential, red (acidic) and blue (basic). The region of Sth1 that sits on top of the nucleosome’s acidic patch is highlighted with a red dashed ellipsoid and a separate interaction with helix 1 of H2B is highlighted with a yellow ellipsoid. **b**, same as panel (b) but with several nucleosome-binding proteins shown in ribbon representation: Sir3 (3TU4.pdb^32^); RCC1 (3MVD.pdb^35^); LANA peptide (1ZLA.pdb^33^); CENP-C peptide (4INM.pdb^34^); and PRC1 (4R8P.pdb^41^). **c**, Close up of the acidic patch and the “arginine anchor” motif. Cryo-EM density from a sharpened, high-resolution map of RSC^SAR^ is shown, with density for the arginine anchor colored blue.

**Fig. 6.**
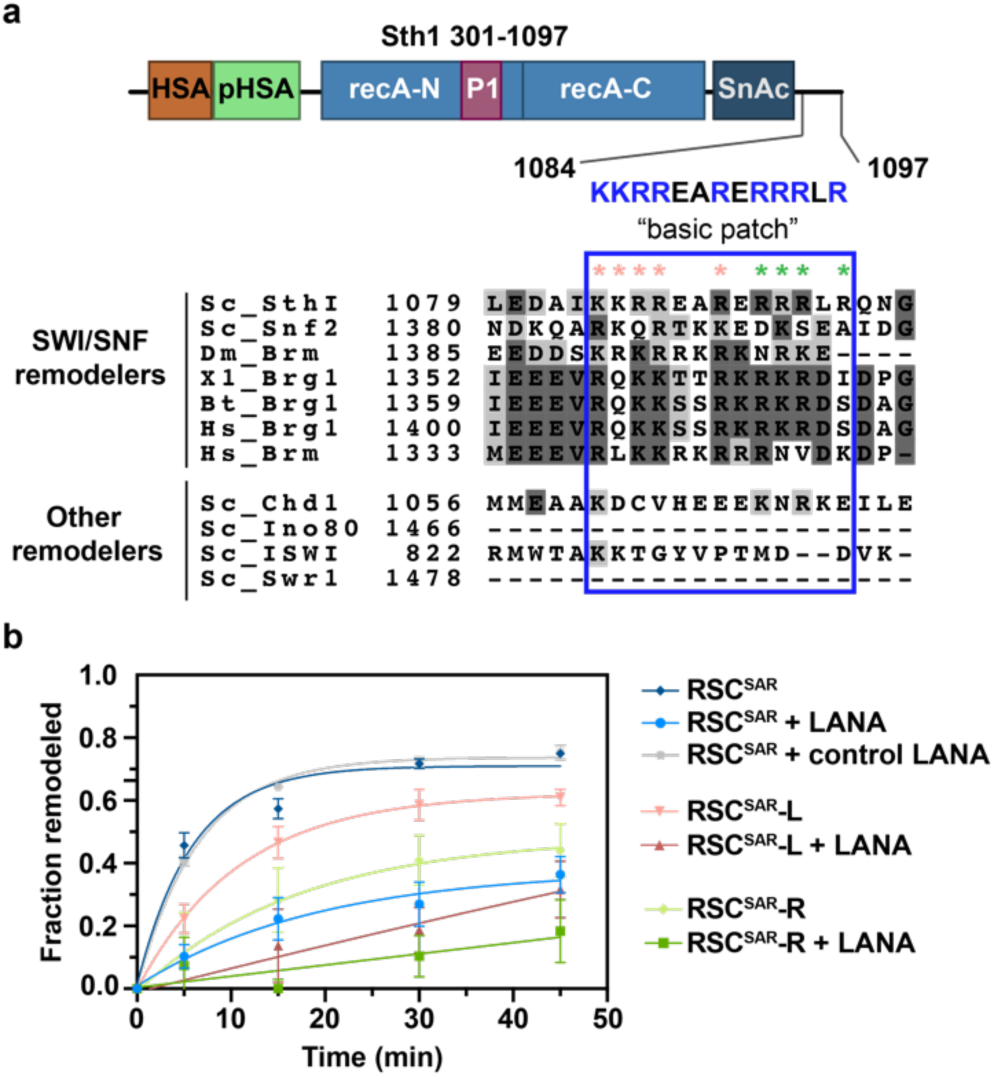
Mutations in a basic region of Sth1 disrupt nucleosome remodelling. **a**, Schematic of the Sth1_301-1097_ construct used in the cryo-EM analysis (top). A “basic patch” at the C-terminus is highlighted, with a sequence alignment shown below. Red and green asterisks indicate residues mutated in (b). **b**, Restriction enzyme accessibility assay of wildtype and mutant RSC^SAR^ complexes. The LANA peptide binds the acidic patch and has been shown to reduce remodeling rates in other systems^42^. Error is reported as standard deviation and curves were fit to a one-phase association non-linear regression curve. R and L mutants correspond to residues highlighted in panel (a) with pink (L) and green (R) asterisks.

Based on our cryo-EM maps and sequence constraints, we hypothesized that SWI/SNF remodelers use the basic patch to bind the nucleosome acidic patch and that this interaction may promote remodeling. To test this, we generated two RSC^SAR^ mutants where the basic residues in either the first half (RSC^SAR^-L) or second half (RSC^SAR^-R) of the patch were mutated to alanine (Fig. 6a, colored asterisks). Remodeling was monitored using a restriction enzyme accessibility assay. In agreement with our hypothesis, basic patch mutants showed a decrease in remodeling compared to wildtype (Fig. 6b, Supp. Fig. 9). To further test whether the reduction in remodeling was due to a compromised interaction between the nucleosome’s acidic patch and Sth1’s basic patch, we performed a competition assay. We pre-incubated nucleosomes with the LANA peptide from Kaposi’s sarcoma herpesvirus^32–35^, a known acidic patch binder, and assayed for remodeling activity. Both wildtype and mutant RSC^SAR^ complexes showed a significant decrease in remodeling in the presence of the LANA peptide, but were unaffected by a LANA control (where basic residues are mutated to Ala) (Fig. 6b). In addition to the cryo-EM density observed immediately above the acidic patch, we noticed another, stronger density along the surface of helix 1 of H2B (Fig. 5a, yellow dashed circle and Supp Fig 8a,b). The region of Sth1 C-terminal to the SnAc domains appears to extend along the surface of the nucleosome and contacts both the acidic patch and helix1 of H2B. This interaction with H2B seems to be unique to SWI/SNF remodelers, as other known acidic patch interactors (LANA peptide, RCC1, Sir3, CENP-C, PRC1) do not colocalize to this region (Fig. 5b,c). The resolution of our cryo-EM reconstruction is not sufficient to build a molecular model for any of the density that engages the surface of the nucleosome, and further analysis will be required to better understand the molecular detail of this interaction.

## Discussion

In this work, we have used a well-studied RSC sub-complex (here termed “RSC^SAR^”) to understand, mechanistically, how the essential ARP subunits regulate its remodeling activity. We determined a cryo-EM reconstruction of the RSC^SAR^ complex, defining its overall architecture (Fig. 1) and the conformation of a regulatory hub within the ATPase domain (Fig. 2). Our structure and biochemical analysis showed that binding of the ARPs induces a helical fold in the HSA domain of Sth1 (Fig. 3), and comparisons between our structure of RSC^SAR^ and those of the related Snf2 ATPase bound to a nucleosome (in the absence of the ARPs) revealed that the folded HSA domain forms a pseudo-helical bundle with two structural elements of the regulatory hub: the post-HSA domain and P1 domains (Fig. 2). The most striking overall feature of the RSC^SAR^ structure is the position and flexibility of the ARP module. In RSC^SAR^, the module lies on the same plane as the nucleosome and adopts a large number of conformations about a pivot point located where the HSA helix meets the post-HSA/P1 regulatory hub (Fig. 1c and Supp. Fig. 3d). In the RSC complex, in contrast, the ARP module is rotated by ∼120°, bridging the ATPase to the Substrate Recognition Complex (SRC), which interacts with the opposite face of the nucleosome (Fig. 1d, e). Despite these large conformational differences, one aspect of the ARPs remains constant: they do not interact with anything other than the HSA helix in RSC^SAR^, full RSC, or the related yeast SWI/SNF complex. Taking all of this together, we propose a two-step model for the ARP regulation of remodeling in RSC (Fig. 7). First, binding of the ARPs to the HSA domain in the ATPase stabilizes its helical fold (Fig. 7a), with the HSA helix in turn interacting with and organizing the other elements in the regulatory hub (Fig. 7b). Second, assembly of the full RSC complex results in a rearrangement of the HSA helix, which now bridges the ATPase, through its contact with the regulatory hub, to the SRC (Fig. 7c). We propose that this connection, whose rigidity (i.e. helical nature) depends on the presence of the ARPs, is responsible for coupling the nucleotide state and conformation of the ATPase to the SRC.

**Fig. 7.**
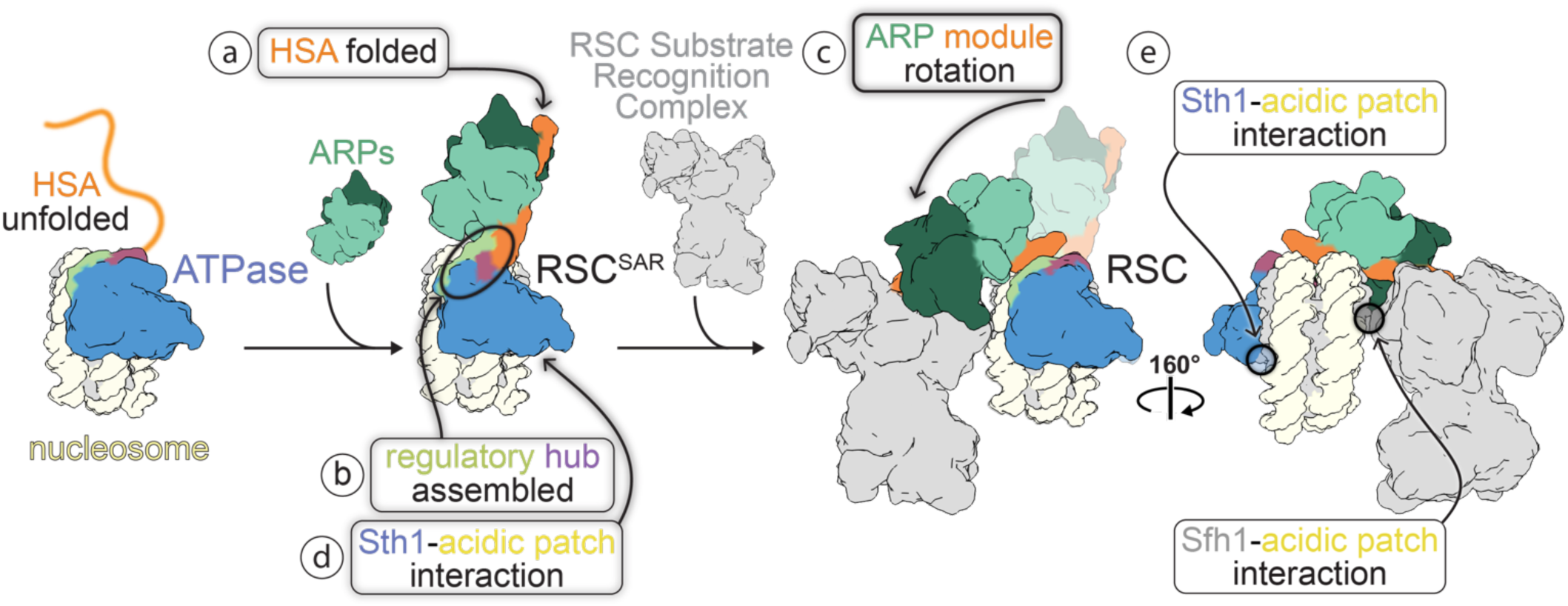
A model for ARP function in chromatin remodeling and RSC assembly. **a-e**, Model for ARP binding and full RSC assembly. ARP module binding induces a helical fold of the HSA domain (a), which in turn leads to the full assembly of the regulatory hub (b). Incorporation into the full RSC results in a large conformational change in the ARP module, which now connects the ATPase domain to the Substrate Recognition Complex (SRC) (c). When assembled, RSC interacts with the acidic patch on both faces of the nucleosome, with these interactions being mediated by Sth1 (d,e) and Sfh1 (e).

The striking difference in the position of the ARP module in RSC^SAR^ vs. RSC may have important implications for RSC assembly. Previous studies have proposed that human SWI/SNF complexes assemble in a hierarchical fashion^36^, with the so-called “ATPase module”, roughly equivalent to the RSC^SAR^ complex used in our study, operating as an independent sub-complex that becomes loaded onto the bulk of the SWI/SNF complex in the final stages of assembly. Our data suggests that not only does the assembly of SWI/SNF complexes require an ordered addition of subunits as suggested^36^, but that the position of the ARP regulatory module must undergo a rearrangement for the full remodeling complex to reach its final, most active form (Fig. 7c). As outlined above, controlling the conformation of the HSA helix is likely central to the function of the remodeler and may synthesize inputs from the SRC module and the ATPase itself. As SWI/SNF complexes are both conformationally and compositionally heterogeneous, there are likely additional stages along the assembly pathway. Further analysis, particularly of assembly intermediates, will be necessary to evaluate this possibility.

Although our structure and those of the full SWI/SNF complexes from yeast, SWI/SNF and RSC, showed the ARPs interacting only with the HSA helix, a recent cryo-EM structure of the BAF complex, the human ortholog of SWI/SNF, suggested that the ARP module might associate with the N lobe of the ATPase^24^. Although this may reflect organismal differences, it should be noted that the BAF structure was determined in the absence of nucleotide while the SWI/SNF and RSC structures were determined in the presence of ADP-BeF_3_; it is therefore possible that the conformation and interaction network of the ARP module changes throughout the remodeling cycle as the ATPase changes nucleotide state. A complete, high-resolution structural characterization of a SWI/SNF complex in all nucleotide states remains essential to fully understand the remodeling reaction.

During our processing we found that a subset of particles showed a “peeled” DNA conformation at the nucleosome exit site. Using focused classification, we were able to determine a 4.3 Å cryo-EM structure of the RSC^SAR^-nucleosome complex with an exit site DNA conformation that shows an approximate 40° peeling from the canonical nucleosome DNA conformation (Fig. 4a). The conformation requires breaking of several DNA-histone contacts mediated by basic residues in histones H3 and H2A. Previous reconstructions of Snf2 bound to the nucleosome did not report this peeled DNA conformation^19,29^. A major difference between this study and previous Snf2 cryo-EM studies is the inclusion of the ARP module in our work, suggesting that addition of the ARP module may aid the ATPase in peeling the exit site DNA. Although this observation may also be due to the individual processing pipelines used in each study or reflect differences between Snf2 and Sth1, mutations in the Snf2 N-lobe near the exit site DNA severely reduce the ability of the isolated enzyme to remodel nucleosomes, supporting the idea that peeling is an intrinsic property of the ATPase domain^19^. Since other RSC subunits have also been shown to interact with the exit site DNA^21,22^, it is likely that multiple components of the full complex act in this process.

Lastly, our cryo-EM analysis of RSC^SAR^ showed that the ATPase interacts with the surface of the nucleosome, in agreement with a recent structure of RSC bound to the nucleosome^21^. Our cryo-EM and biochemical analysis allowed us to identify a small, highly basic region of Sth1 that interacts with the nucleosome’s acidic patch, a hotspot to which many remodelers have been previously shown to bind (Fig. 7d). For example, CHD1 is reported to interact with the acidic patch^37^, but lacks the C-terminal basic motif identified in this work, suggesting that it uses alternative mechanisms to those employed by SWI/SNF remodelers. Importantly, the human ortholog of Sth1, Brg1, was shown to have decreased remodeling of nucleosomes carrying acidic patch mutations^38^. Based on sequence constraints from our Sth1 construct, we propose that a stretch of basic residues C-terminal to the ATPase domain, which is conserved between SWI/SNF family members (Sth1 and Snf2 from yeast) and throughout evolution (yeast, human, and fly orthologs), directly binds to the acidic patch (Fig. 6a). A recent study showed that ISWI also contains a small basic stretch C-terminal to the ATPase domain that mediates binding to the acidic patch ^39^. This motif is conserved in ISWI orthologs and mutating it reduces the rate of remodeling, as we have reported here. This motif in ISWI seems to have a strong interplay with accessory domains in ISWI that do not exist in SWI/SNF remodelers, so further analysis of the acidic patch interaction in the context of each remodeler is necessary to understand if and how it contributes to the functional diversity of remodelers. Intriguingly, we observe an additional interaction along the H2B surface that is distinct from the interaction with the acidic patch (Fig. 5a, yellow circle). Several studies have recently reported that SWI/SNF remodelers engage the acidic patch on the opposite face of the nucleosome using their Sfh1 (RSC) or Snf5 (SWI/SNF) subunit (Fig 7e)^20–22,24,24^. This interaction is vital to the function of the enzyme, as mutations in a conserved basic domain in SMARCB1 (Snf5 homolog) disrupt binding to the nucleosome, reduce remodeling rates, and have defects in cell-type differentiation^40^. It therefore appears that RSC, and likely SWI/SNF remodelers more broadly, is able to simultaneously engage the acidic patch on both faces of the nucleosome (Fig. 7e). This two-sided engagement with the histone octamer is likely a key component of the remodeling reaction and would allow RSC to maintain hold of the histone octamer as it threads DNA around the nucleosome.

## Materials and Methods

### Sample preparation and biochemical analysis

#### Protein expression and purification

Sth1 constructs were cloned as described^41^. Sth1 constructs were expressed in BL21 RIPL cells and grown in TB supplemented with kanamycin and chloramphenicol until reaching an OD_600_ of ∼0.8. Cells were induced with 0.5 m M isopropyl β-D-1-thiogalactopyranoside (IPTG) at 16 °C and grown overnight, followed by centrifugation. Cell pellets were resuspended in lysis buffer (20 mM Tris pH 8, 300 mM NaCl, 2 mM imidazole, 1 mM DTT, and 10% glycerol) supplemented with protease inhibitors and DNase, lysed by sonication, and clarified by centrifugation. Lysates were applied to tandem 1 mL HiTrap FF columns (Ge Healthcare) charged with Ni^2+^, and washed with 20 mM imidazole. Proteins were eluted with a gradient of 20-250 mM imidazole. Proteins were diluted to 75 mM NaCl and loaded onto a MonoQ 5/50 GL (GE Healthcare) equilibrated in QO buffer (20 mM Tris pH 8, 50 mM NaCl, 1 mM DTT, 5% glycerol). Proteins were eluted with a 50-600 mM gradient, and then run over a final S200 10/300 Increase (Ge Healthcare) size exclusion column equilibrated in 20 mM Tris pH 7.5, 150 mM NaCl, 1 mM DTT, and 5 % glycerol. For assembly of ATPase module complexes, Sth1 was mixed with Arp7/Arp9/Rtt102 in a 1:2 ratio and incubated on ice for a minimum of 30 minutes prior to use.

*Xenopus laevis* histones were expressed in BL21-Rosetta cells and purified from inclusion bodies, essentially as described in^42^. DNA containing the Widom 601 positioning sequence was purified using restriction enzyme digestion of a plasmid containing 16 copies of the 601 sequence ^43^. Nucleosomes were assembled using the salt-gradient dialysis method, essentially as described^44^.

#### GraFix crosslinking

Samples were crosslinked essentially as described^45^. Briefly, nucleosomes (185 bp DNA, *X. laevis* histones) and yeast RSC^SAR^ (Sth1_301-1097_, Arp7, Arp9, Rtt102) were combined at a final concentration of 3 μM and 9 µM, respectively, in a base buffer containing 20 mM HEPES, pH 7.5, 50 mM KOAc, 3 mM MgCl_2_, 1 mM ADP, 1 mM BeF_3_. Glycerol gradients were made by layering 10 mL of light solution (base buffer + 10% glycerol) on top of 2 mL of heavy solution (base buffer + 30% glycerol + 0.10% glutaraldehyde) and mixing the solution using a BioComp Gradient Master 107 set to rotate for 90 seconds with an 83° tilt. Gradients were allowed to settle at 4°C for at least 1 hour before centrifugation. 100 μL of protein solution was layered on top of the glycerol gradient and spun using an SW-60 rotor at 100,000 g for 18 hours at 4°C. Samples were fractionated and analyzed using native PAGE. Fractions containing cross-linked RSC^SAR^-nucleosome samples were pooled, dialyzed in base buffer to remove glycerol, and concentrated to ∼5 μM.

#### Remodeling assays

In a 40 μL reaction, proteins at 10 nM were incubated for 5 minutes with 20 nM 225 bp nucleosomes (EpiCypher) at 30 °C in 20 mM Tris pH 7.5, 50 mM potassium acetate, 2 mM MgCl_2_, 0.1 mg/mL BSA, 1 mM DTT, 2 units MfeI, and 2 units PmlI. The remodeling reaction was started with the addition of 1 mM ATP. 5 uL aliquots were removed at each time point and added to 2 uL 5 mg/mL Proteinase K (NEB) in 5% SDS, and incubated at 50°C for 30 minutes. Samples were analyzed in quadruplicate on 8-16% TGX gels (BioRad) that had previously been equilibrated in 0.5X TBE for 30 minutes at 150V. Gels were stained with Sypro ruby (Thermofisher) and quantified by densitometry using the program ImageLab (BioRad). Each sample, the 225 bp DNA band was quantified and normalized relative to the no-protein control to obtain the fraction of remodeled nucleosome.

### Cryo-EM structure determination and model building

#### Sample preparation and data collection

All grids were made following the same general protocol. UltrAuFoil R1.2/1.3 300 mesh grids (Quantifoil GmbH) grids were glow-discharged for 30 seconds at 25 mAmp and used within 15 minutes. 4 μl of sample was applied to a charged grid and blotted and plunge-frozen in nitrogen-cooled liquid ethane using a Vitrobot Mark IV robot (Thermo Fisher) set to blot force 20, blot time 4 s, 100% humidity, 4° C. Grids were made using 1 µM - 5 µM sample, with concentrations increased 2-5 fold when supplemented with 0.05% n-Octyl-β-D-Glucopyranoside (β-OG).

Data were collected using a Talos Arctica 200 keV TEM (Thermo Fisher) operating in nanoprobe mode and equipped with a K2 Summit Direct Detector (Gatan) operating in counting mode. For the RSC^SAR^ apo sample (Arp module), a Volta Phase Plate was used. The VPP was advanced every 30 minutes to ensure phase shifts < 135°. Magnification was set to 36,000 for a final pixel size of 1.16 Å and defocus was set to -0.6 μm to -2.5 μm. Exposure rate was 5-7 e^-^/pixel/s, depending on dataset, with 200 ms frames, and a total exposure of 50-55 e^-^/ Å^2^. New camera gain references were collected before each dataset and the hardware dark reference was updated at least every 24 hours. Parallel illumination of the microscope was performed according to^46^. Holes containing thin ice were manually queued for automatic data collection using the Leginon software suite. Data were processed on-the-fly using the Appion software suite^47^ and each micrograph was visually inspected. Subpar micrographs (ethane contamination, crystalline ice, empty holes, etc.) were manually excluded from the dataset.

#### Arp module cryo-EM structure determination

RSC^SAR^ was purified by recombinant expression in E. coli (see above). Grids were prepared as described, with a range of protein and detergent concentrations. A total of 5 datasets were combined and processed as a single dataset. All datasets were collected with similar exposure rates, total exposure, frame rate, and magnification, and processed as a single dataset. A total of 5365 movies were collected. MRC stacks were compressed to TIF using the mrc2tif script in IMOD^48^. Gain correction, motion correction, dose weighting, and CTF estimation were performed using MotionCor2^49^ and CTFFIND4^50^ in Relion-3^51^. Micrographs with defocus values > -2.5 μm and CTF fits worse than 5 Å were removed from the dataset. Particles were picked from motion-corrected micrographs using crYOLO^52^. A total of 1,986,341 particles were picked and subjected to multiple rounds of 2D classification. An initial model was determined using cryoSPARC^53^, yielding a structure that contains clear density for the Arp module (Sth1 HSA helix, full-length Arp7/Arp9/Rtt102). The Sth1 ATPase domain is disordered in our map, and extensive effort to find a well-ordered population did not yield a structure. 3D classification in Relion-3 was used to find a subset of 415,957 particles that classified into models with well-defined secondary structure. 3D refinement yielded a structure with a GSFSC resolution of ∼4.6 Å resolution (unmasked). The resolution improved to ∼4.2 Å after masking and post-processing in Relion-3. The unsharpened, sharpened, and half maps were deposited in the EM Data Bank as EMD-21489.

#### Arp module model building and validation

Our cryo-EM map shows clear density for the RSC Arp module, including the Arp7/Arp9/Rtt102 heterotrimer, and the HSA helix of Sth1. To build a molecular model of the RSC Arp module, the crystal structure of Snf2_HSA_/Arp7/Arp9/Rtt102 (4I6M.pdb^54^) was docked into the map. The Snf2 HSA helix was mutated to Sth1 identity and numbering and manually rebuilt in certain regions using Coot^55^. After docking, we realized that there was density in the ATP-binding pocket of Arp7, despite no ATP being included in our buffers. This site is enzymatically dead and the ligand is likely from the recombinant expression. To model ATP in this binding site, the ATP ligand from the Arp7_ATP_/Arp9/Rtt102 crystal structure (5TGC.pdb^41^) was docked into the map. To refine our model, we used a cloud-based Rosetta pipeline^56^. Initially, ∼1000 models were generated using RosettaCM^57^. The top 10% based on Rosetta energy score were scored using MolProbity^58^. The ten best models based on MolProbity score were refined using Rosetta Relax, which improved the MolProbity and Clashscore for all models. A final model consisting of 10 models was deposited as 6VZG.

#### RSC^SAR^-nucleosome cryoEM structure determination

RSC^SAR^ -nucleosome was purified and crosslinked as described above. Multiple data sets were collected on grids with various protein and detergent concentrations. All datasets were collected with similar exposure rates, total exposure, frame rate, and magnification, and processed as a single dataset. A total of 9292 movies were collected. MRC stacks were compressed to TIF using the mrc2tif script in IMOD ^48^. Gain correction, motion correction, dose weighting, and CTF estimation were performed using MotionCor2^49^ and CTFFIND4^50^ in Relion-3^51^. Micrographs with defocus values > -2.5 μm and CTF fits worse than 4 Å were removed from the dataset. Particles were picked from motion-corrected micrographs using crYOLO^52^. A total of 2,020,734 particles were picked and initial 2D classification showed structures that were clearly nucleosomes with density attached to SHL2 and SHL-2. An initial model was determined using cryoSPARC^53^, yielding a 3D structure with clear density for the nucleosome and either one or two copies of Sth1. Additional fragmented density was seen at the N-terminus of Sth1. Instead of using 2D classification to ‘clean’ our dataset, we used multiple rounds of 3D classification to isolate particles that partitioned into clear nucleosome-bound structures. This yielded a ‘clean’ dataset of 747,408 particles.

Refinement of the ‘clean’ dataset yielded a reconstruction at 3.7 Å resolution after masking and post-processing. The full dataset clearly has a mixture of particles with either 1 or 2 copies of Sth1 bound, yielding a structure with full occupancy at SHL2 and partial occupancy at SHL-2. To improve the density of the Arp module, which shows fragmented density in this reconstruction, we used 3D classification. A total of 293,940 particles partition into 3D classes with improved Arp density, and refine to a final resolution of ∼3.9 Å after masking and post-processing. To further improve the Arp density, multi-body refinement^59^ was performed using masks for 3 bodies (nucleosome, Sth1, Arp module). Masks were partially overlapping and included > 100 kDa of mass for each body. This dramatically improved the density of the Arp module, and yielded moderate improvements in resolution for the nucleosome and Sth1 ATPase domain. Unsharpened, sharpened, and half-maps for the consensus refinement and the three bodies of the multi-body refinement were deposited in the EMDB as EMD-21484.

#### RSC^SAR^ -nucleosome model building and validation

After multi-body refinement, our cryo-EM map shows clear density for the nucleosome, the ATPase domain of Sth1 bound to SHL2, and the Arp module bound to the HSA domain of Sth1. To build the molecular model of RSC^SAR^ bound to the nucleosome, we first docked in the model of a related structure, yeast Snf2 bound to the nucleosome in the presence of Mg-ADP-BeF_3_ (5Z3U.pdb^29^). The Snf2 model was mutated to Sth1 residue number and sequence identity in Coot. In addition to the ATPase domain, clear density is seen for the HSA domain and the two lobes of the Arp module, allowing for un-ambiguous docking of our RSC Arp module structure (Sth1 HSA, Arp7/Arp9/Rtt102) into the map. After rigid-body docking these two models, the main structural differences were in the HSA, post-HSA, and protrusion domains of Sth1. Notably, the HSA and post-HSA domains are more ordered in our map compared to the starting models. The helices were extended with ideal geometry in Coot. However, at this point the termini of these two domains were incompatible (the HSA domain ended at residue 384 and the post-HSA domain began at 381). To relieve this clash, we instead modeled the post-HSA and protrusion domains from another Snf2 structure into our map (5HZR.pdb^60^). This yielded a post-HSA conformation with much better agreement relative to the position of the HSA domain. To mitigate bias in our modeling, we created multiple hybrid models using a variety of conformations of the post-HSA and protrusion domains of 5HZR.pdb. The position of the post-HSA and protrusion domains can vary significantly depending on how the automatic or manual docking is performed, so we made a variety of models and used these as inputs for RosettaCM^57^. Approximately 1000 models were generated and ranked using Rosetta energy score^61^ and MolProbity score^58^. The top ten models from this analysis had good agreement in the post-HSA domain. The nucleosome was refined using phenix.real_space_refine. The top ten Sth1 models were combined with the real space refined nucleosome model and further refined using Rosetta Relax, which improved geometry and clashscores for all models. The nucleosome DNA was replaced with the refined coordinates from phenix.real_space_refine^62^. The Arp module was not refined in the map, but rather rigid-body docked and slight modifications to Sth1 residues 362-383 were made in Coot^55^. Lastly, B-factors were refined in Phenix^63^. A final model consisting of 10 models was deposited in the PDB as 6VZ4.

#### RSC^SAR^-nucleosome “peeled DNA exit site” cryoEM structure determination

During processing we noticed that 3D classification often showed heterogeneity in the DNA exit site of the nucleosome. To better classify this heterogeneity, we used signal subtraction and 3D classification without alignment to sort the 293,940 particles from our consensus refinement based on the conformation of the DNA exit site (Supp. Fig 1). We found the ∼25% of particles segregated into classes that showed a “peeled” DNA conformation at the exit site. A subset of these particles, corresponding to a single 3D class containing 112,364 particles, were re-centered and re-extracted and subjected to gold-standard 3D refinement in Relion3. This yielded a final reconstruction with a resolution of 4.3 Å after masking and post-processing. The unsharpened, sharpened, and half-maps were deposited in the EMDB as EMD-21493.

## Acknowledgements

We thank the UC San Diego Cryo-Electron Microscopy Facility, which was supported in part by NIH grants to Dr. Timothy S. Baker and a gift from the Agouron Institute to UCSD and the UC San Diego Physics Computing Facility for IT support. RWB is a Damon Runyon fellow supported by the Damon Runyon Cancer Research Foundation (DRG-2285-16). JMR is a Merck fellow of the Damon Runyon Cancer Research Foundation (DRG-2370-19). This work was funded by National Institutes of Health Grants R01 GM092895 (AEL) and R01 GM073791 (RD).

## Data Availability

The cryo-EM maps for apo RSC^SAR^ (Sth1_HSA_-Arp7-Arp9-Rtt102 model) are deposited in the EMDB as EMD-21489 and molecular models are deposited in the PDB as 6VZG. The cryo-EM maps for RSC^SAR^ bound to the nucleosome in the presence of ADP BeF_3_ are deposited in the EMDB as EMD-21484 and molecular models are deposited in the PDB as 6VZ4. The cryo-EM maps of RSC^SAR^ bound to the nucleosome in the presence of ADP BeF_3_ with a peeled DNA conformation are deposited in the EMDB as EMD-21493. Unsharpened, sharpened, and half maps were deposited for each EMDB entry.

## Author contributions

RWB, JMR, and PJC performed all the protein production and purification. RWB performed cryo-EM data collection, analysis, and model building. JMR performed nucleosome remodelling assays. TA performed circular dichroism (CD) experiments and analyzed CD data. AEL supervised the structural and biochemical work. All authors participated in writing and editing the manuscript.

## Supplementary Materials

**Supp.Fig.1.**
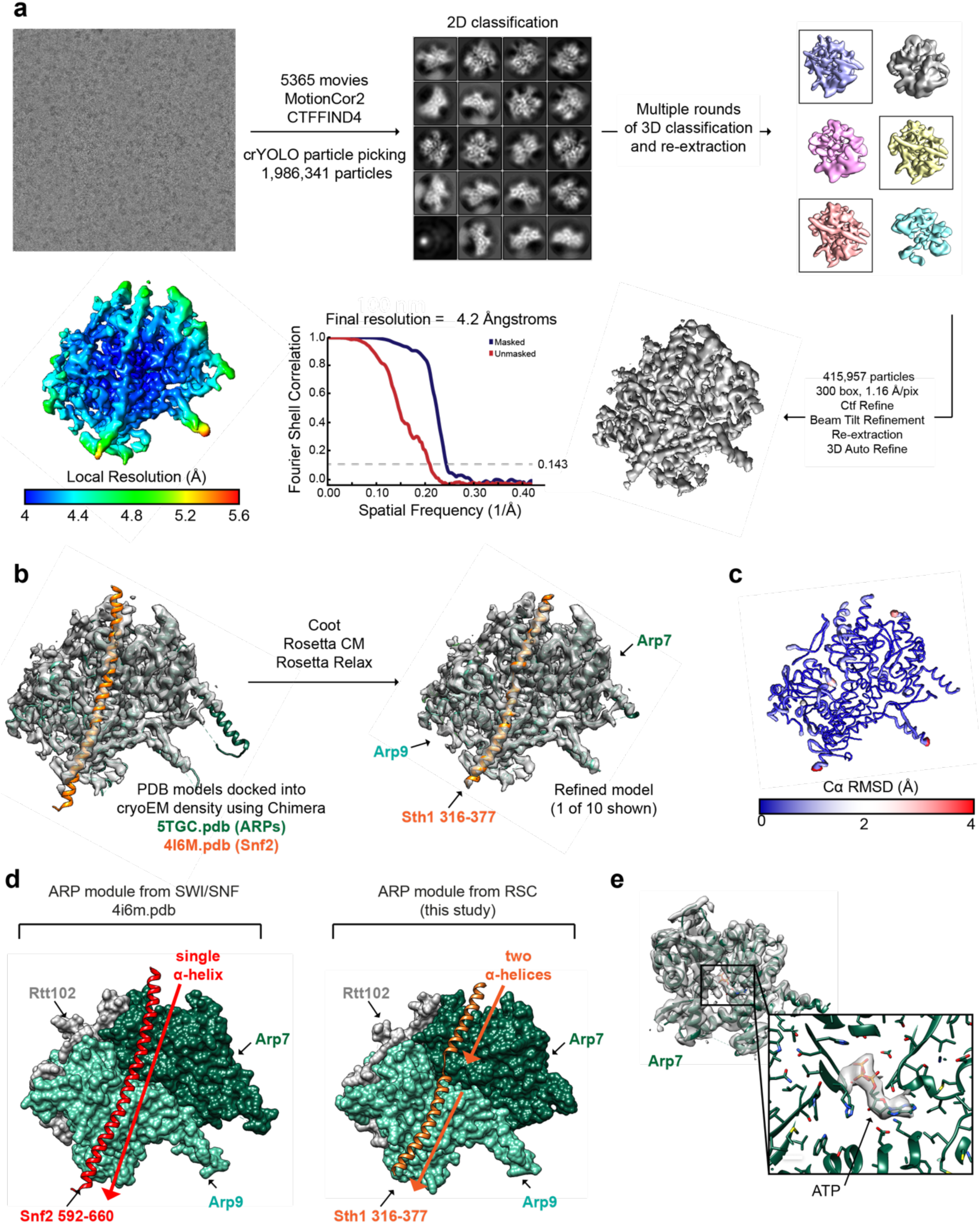
Cryo-EM structure determination of the Sth1-Arp7-Arp9-Rtt102 RSC subcomplex (RSC^SAR^). **a**, Workflow for cryo-EM data acquisition and structure determination. Our processing only revealed density for the ARP module (Sth1_316-377_/Arp7Arp9/Rtt102), accounting for ∼50% of the mass of the complex. An FSC plot and a cryo-EM map colored by local resolution are shown. **b**, Schematic for model building using Coot and Rosetta. Chains from 5TGC.pdb^41^ and 4I6M.pdb^54^ were used as a starting model. **c**, The top 10 Rosetta models were deposited as 6VZG.pdb and the top model is shown with its ribbon thickness and color indicating the RMSD among all top 10 models. **d**, The structure of Snf2_HSA_/Arp7Arp9/Rtt102 (left) has a single, unbroken α-helix for the HSA domain. The structure of Sth1_HSA_/Arp7Arp9/Rtt102 (right) comprises two α-helices separated by a loop. **e**, The cryo-EM map for Sth1HSA/Arp7Arp9/Rtt102 has density for a known ATP binding site in Arp7. This site is catalytically dead^42^ and the ATP is likely from endogenous ATP pools as no ATP was supplemented during purification.

**Supp.Fig.2.**
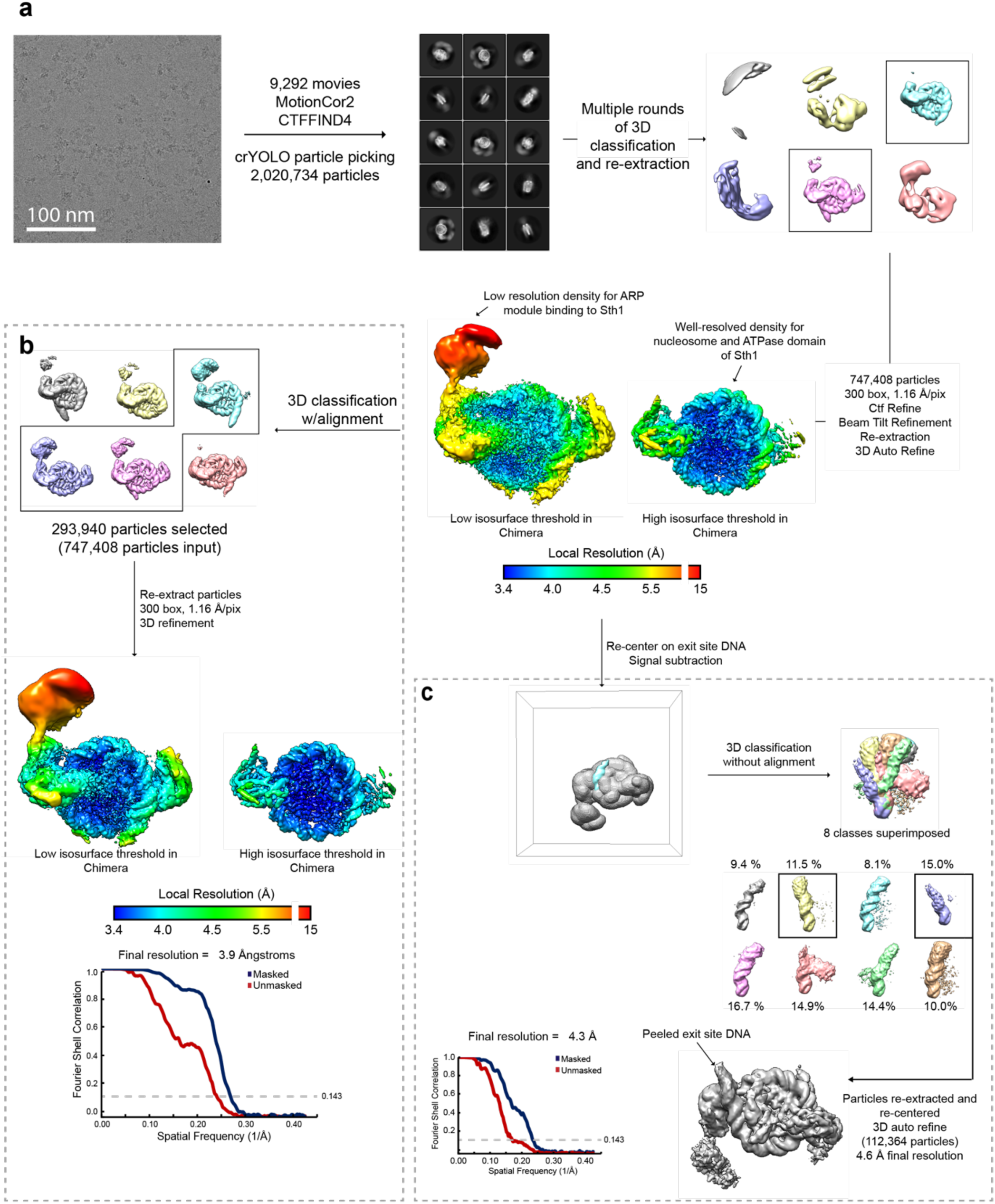
Cryo-EM structure determination of the Sth1-Arp7-Arp9-Rtt102 RSC subcomplex (RSC^SAR^) bound to the nucleosome in the ADP-BeF_3_ state. **a**, Workflow for cryo-EM data acquisition and structure determination. Our processing pipeline yielded a ‘clean’ dataset of ∼750,000 particles that gave a ∼3.7 Å structure, with a local resolution range of 3.4-15 Å. **b**, 3D classification with alignment was used to find ∼300,000 particles with better ARP module density. This dataset reached a final global FSC resolution of ∼3.9 Å with a local resolution range of 3.4-15 Å. **c**, Partial signal subtraction followed by 3D classification without alignment was used to identify a sub-set of particles from our ‘clean’ dataset with an alternate DNA conformation at the nucleosome DNA exit site. A single 3D class comprising ∼110,000 particles was re-centered and re-extracted, yielding a 3D reconstruction with a final overall FSC resolution of 4.6 Å.

**Supp.Fig.3.**
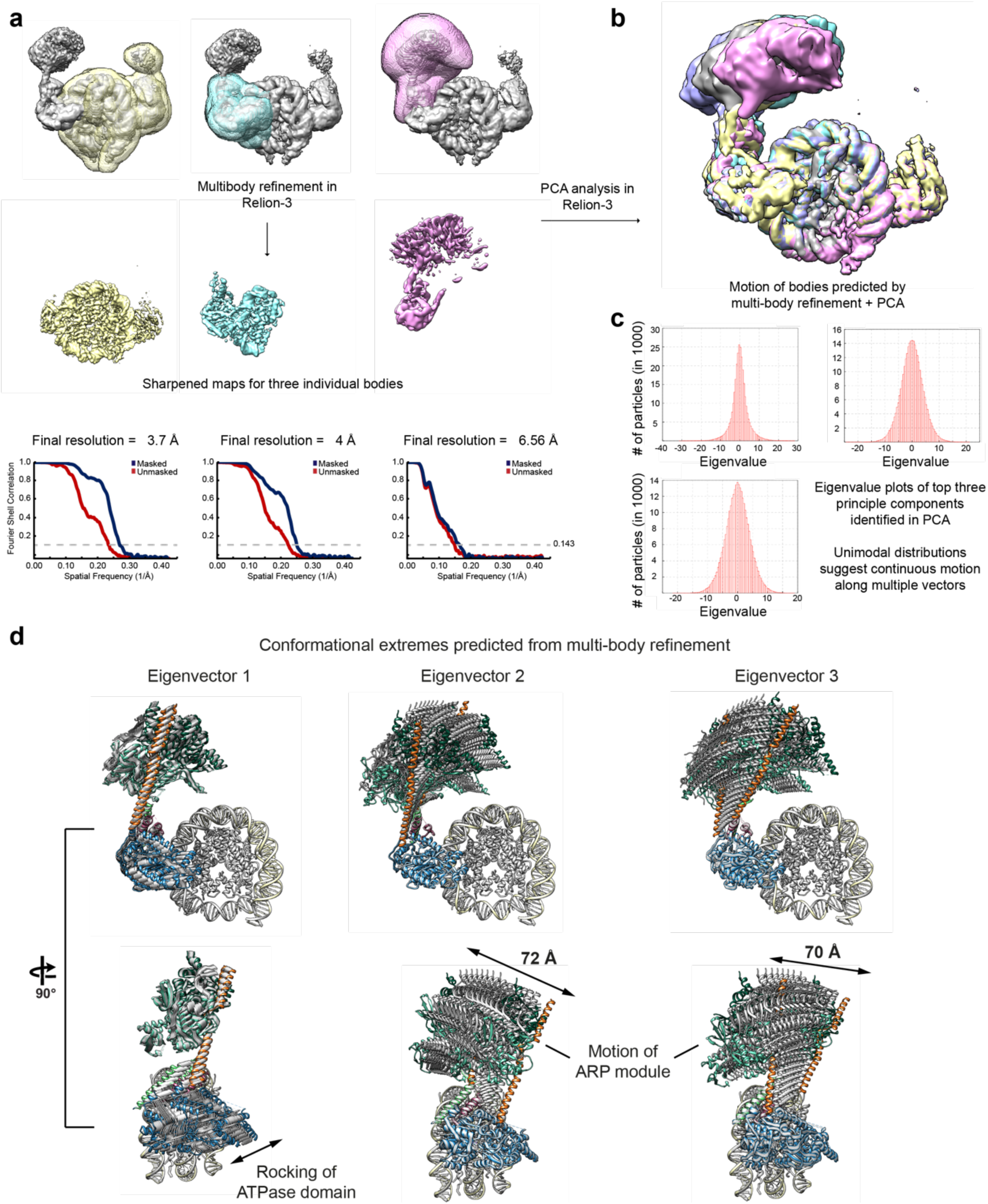
Multi-body refinement of RSC^SAR^ bound to the nucleosome in the ADP-BeF_3_ state. **a**, The consensus refinement of RSC^SAR^_nucleosome from Supp. Fig 2b was used to generate 3 masks and as a starting model for multi-body refinement in Relion-3. This improved the resolution of all regions of the map, especially the ARP module and its connection with the ATPase domain of Sth1. **b**, PCA analysis was performed in Relion-3 and used to generate maps that show the conformational heterogeneity present in our data set. Several of these maps are shown overlaid. **c**, Eigenvalue plots for the top three eigenvectors. **d**, The maps generated in panel b were used to build PDB models. 10 models for each eigenvector were generated and are shown superimposed. The main sources of heterogeneity in our data set are a rocking of the ATPase domain (eigenvector 1) and rocking of the ARP module along two different axes (eigenvector 2 and 3).

**Supp.Fig.4.**
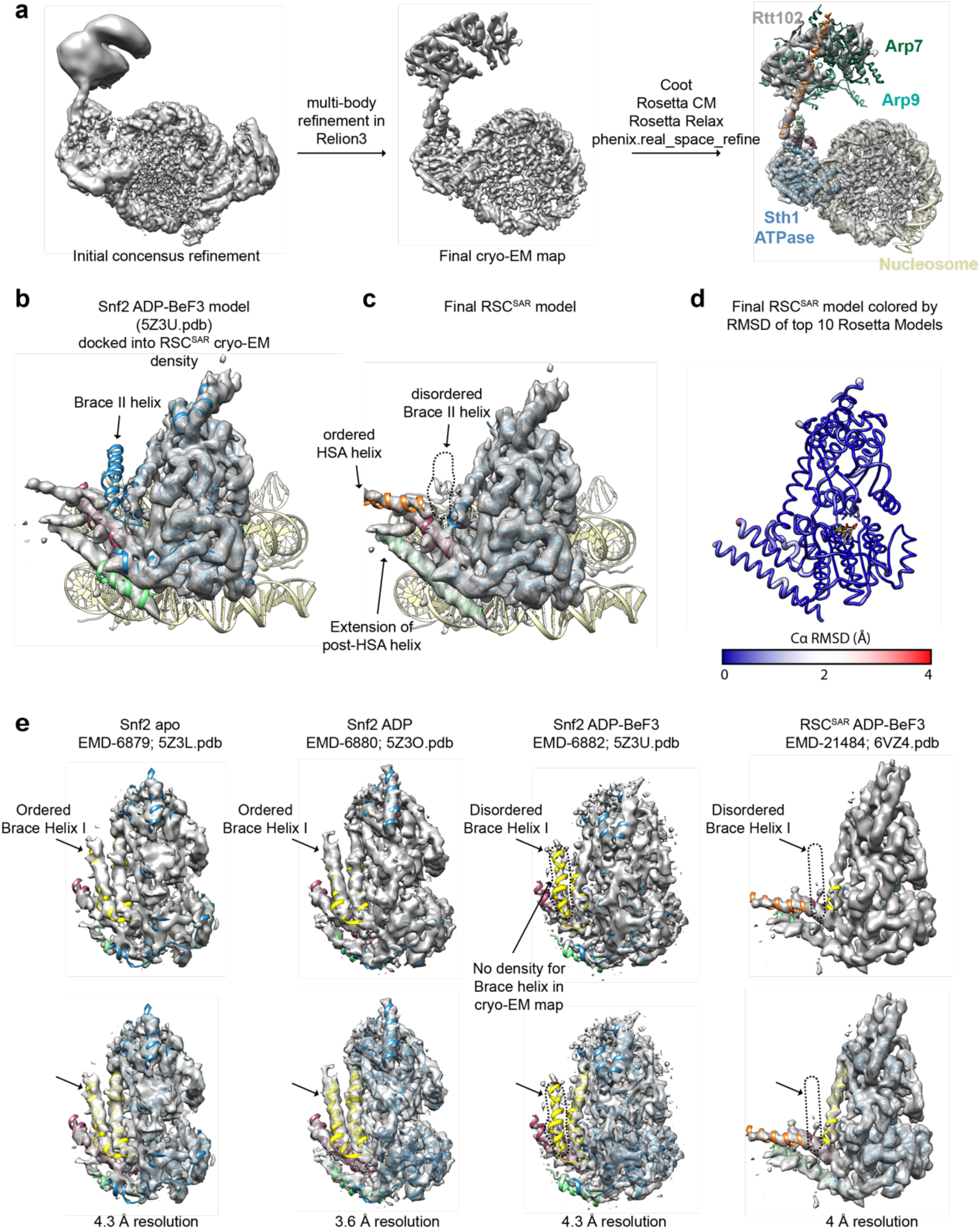
Model building of RSC^SAR^-nucleosome (ADP-BeF_3_) and comparison with Snf2-nucleosome (ADP-BeF_3_). **a**, Model building schematic for RSC^SAR^-nucleosome (ADP-BeF_3_). Using Snf2-nucleosome (ADP-BeF_3_) (5Z3U.pdb^29^) as a starting model, a RSC^SAR^_nucleosome (ADP-BeF_3_) was built and refined using Coot, Rosetta, and phenix.real_space_refine. The ARP module built in Supp. Fig.1 was rigid-body docked into the map, yielding a near-complete model for all components in our sample. **b**, Snf2-nucleosome (ADP-BeF_3_) (5Z3U.pdb^29^) is shown docked into the cryo-EM map of Sth1-nucleosome (ADP-BeF_3_). **c**, the same view as in panel (b) but showing Sth1-nucleosome (ADP-BeF_3_). Structural changes are highlighted. **d**, The top Rosetta model for the ATPase is shown with its ribbon thickness and color indicating the RMSD among the top 10 models. **e**, comparison of all cryo-EM structures of Snf2-nucleosome in the apo, ADP, and ADP-BeF_3_ states and RSC^SAR^-nucleosome (ADP-BeF_3_). Each is shown with cryo-EM map and corresponding PDB model. The same map at the same contour level is shown opaque (top) and transparent (bottom). Note that density for the Brace Helix I is absent in the ADP-BeF_3_ state but present in the apo and ADP-bound states.

**Supp.Fig.5.**
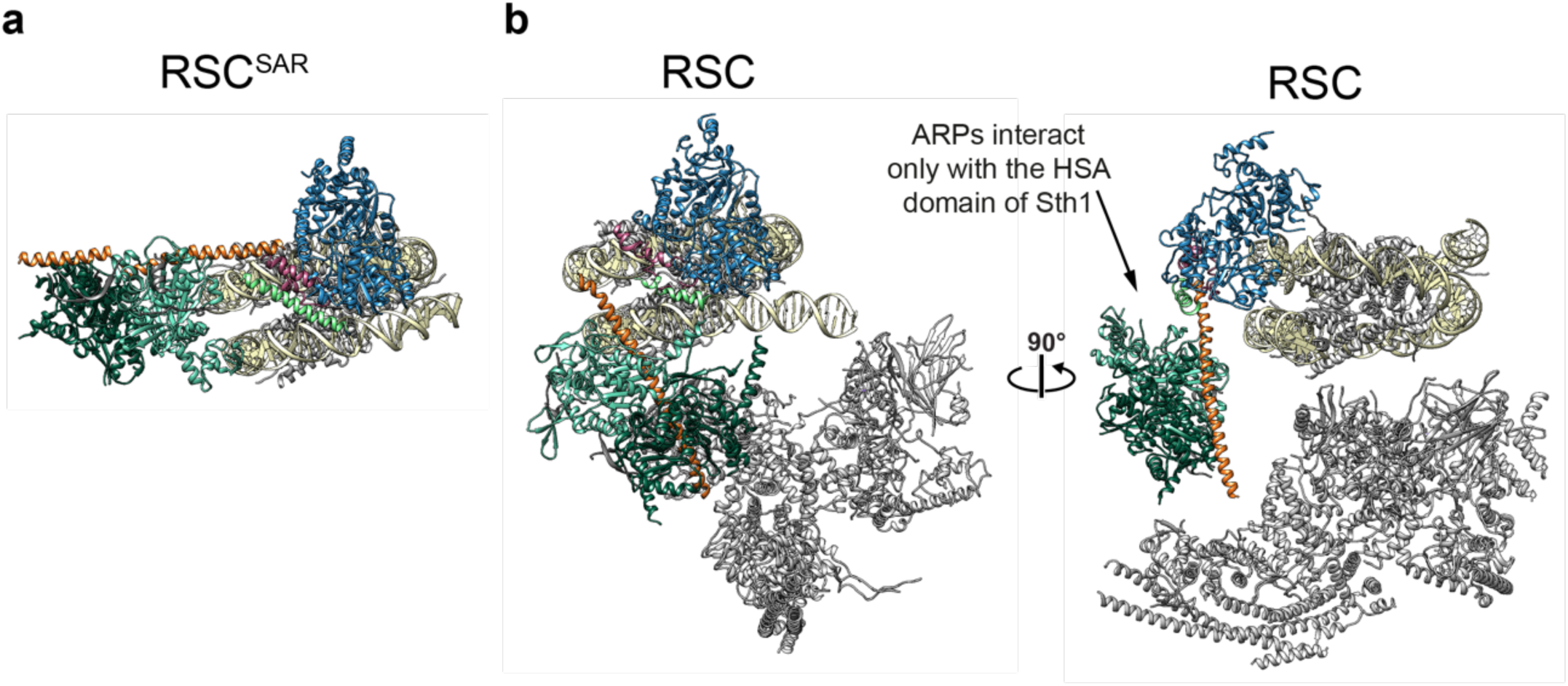
The ARP module of RSC only interacts with the HSA domain of Sth1. **a**, Our model of RSC^SAR^-nucleosome (ADP-BeF_3_). **b**, Model of RSC-nucleosome (6KW3.pdb^22^) is shown in the same orientation as panel (a) (left) and rotated 90° (right). The ARP module only interacts with the HSA domain in both RSC and RSC^SAR^.

**Supp.Fig.6.**
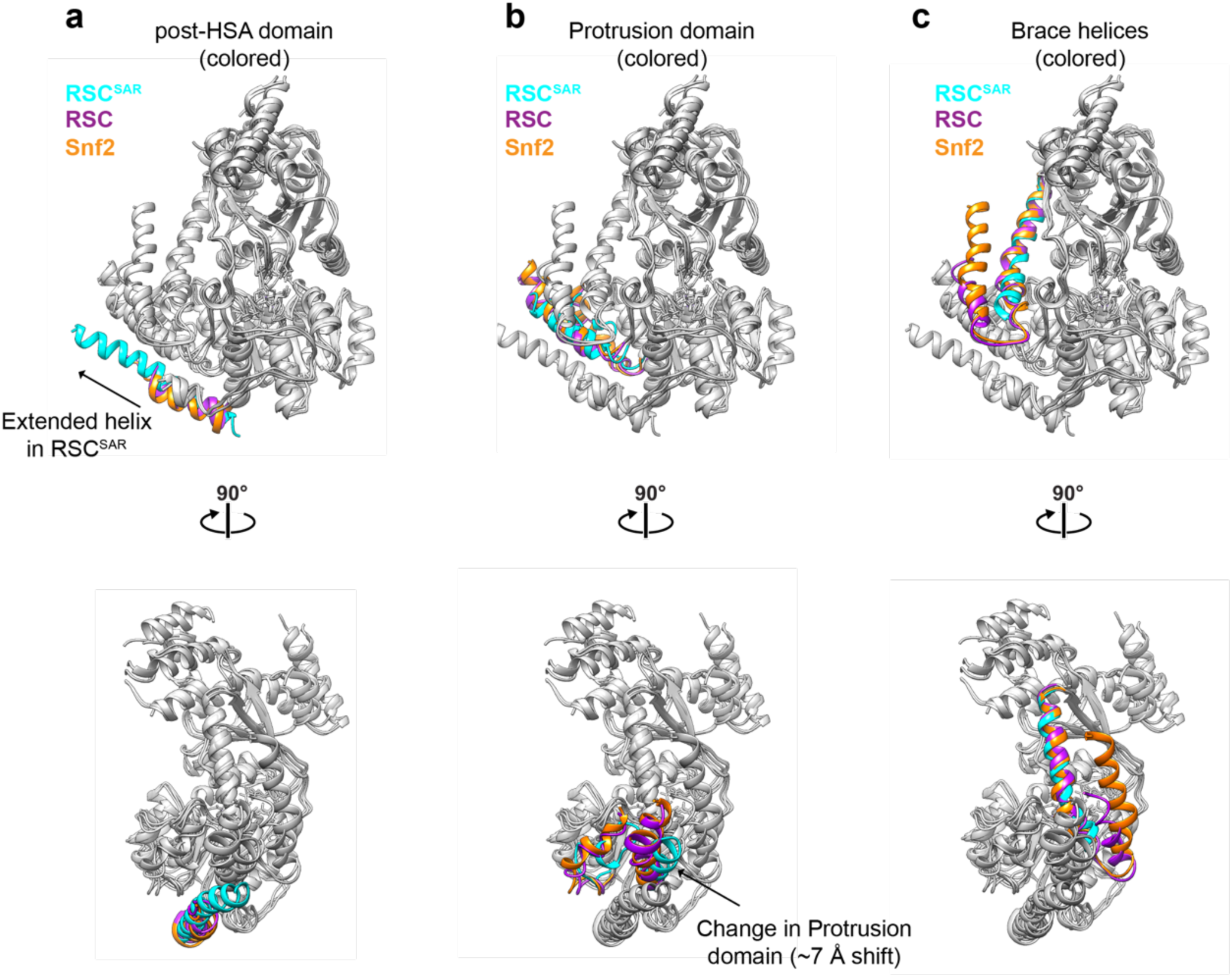
Comparison of the nucleosome-bound ATPase domain of Snf2, RSC, and RSC^SAR^. The PDB model for each ATPase domain was docked into the cryo-EM map of RSC^SAR^. The post-HSA domain (a), Protrusion 1 domain (b), and Brace Helices (c) were colored according to their respective PDB model in each panel. Cyan: RSC^SAR^, Magenta: RSC (6KW3.pdb^22^), Orange: Snf2 (5Z3U.pdb^29^).

**Supp.Fig.7.**
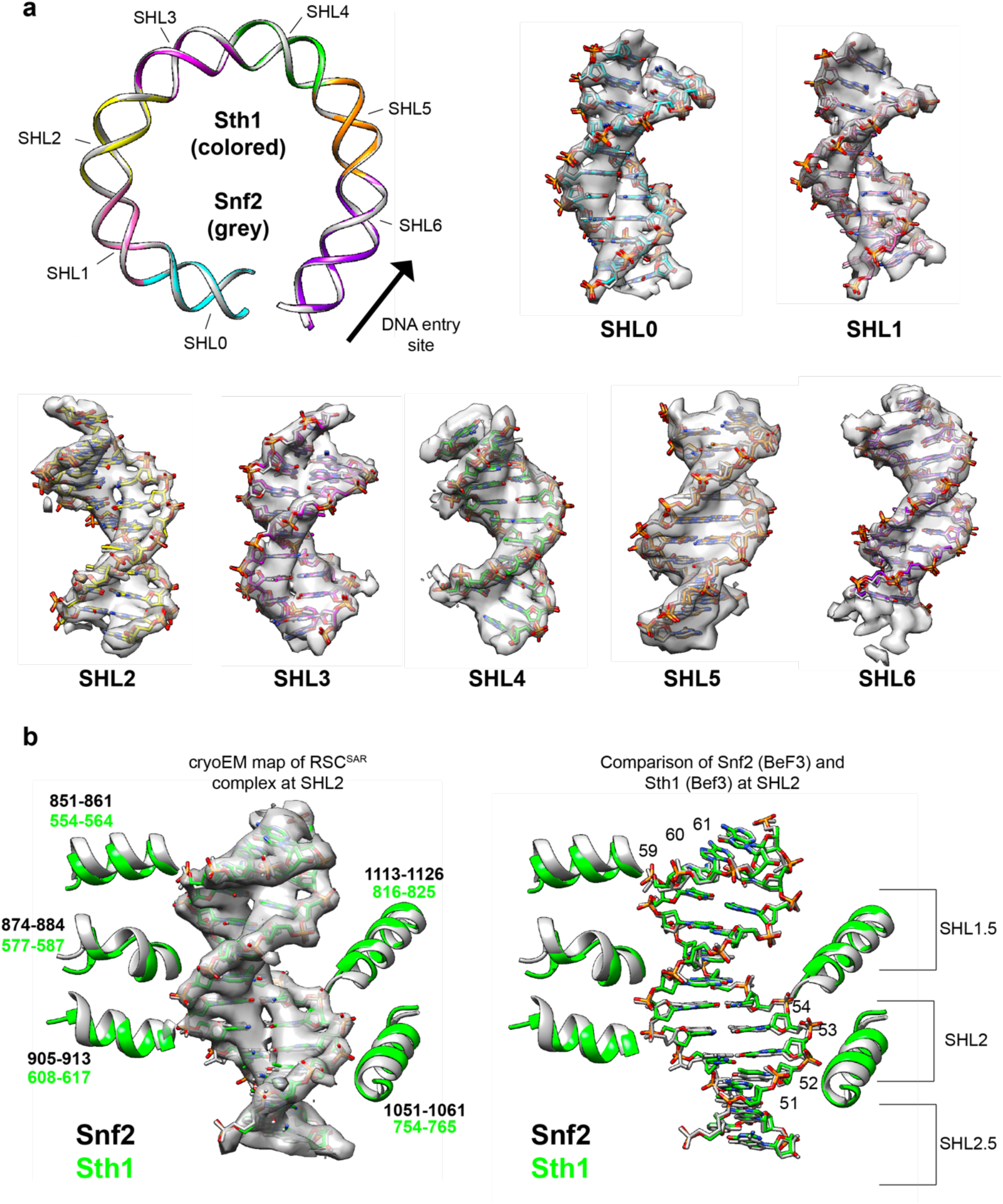
Nucleosome DNA conformation of the RSC^SAR^-nucleosome (ADP-BeF_3_) cryo-EM structure. **a**, The models of Snf2-nucleosome (ADP-BeF_3_) (5Z3U.pdb^29^) and RSC^SAR^-nucleosome (ADP-BeF_3_) were aligned. A ribbon representation of the nucleosomal DNA is shown for Snf2 (grey) and Sth1 (colored by SHL). SHL0 – SHL6 are shown for both models inside the cryo-EM density of RSC^SAR^-nucleosome (ADP-BeF_3_). **b**, SHL2 for both models is shown, including regions of the ATPase that engage the DNA and have been shown to be important for remodeling^19,29^.

**Supp.Fig.8.**
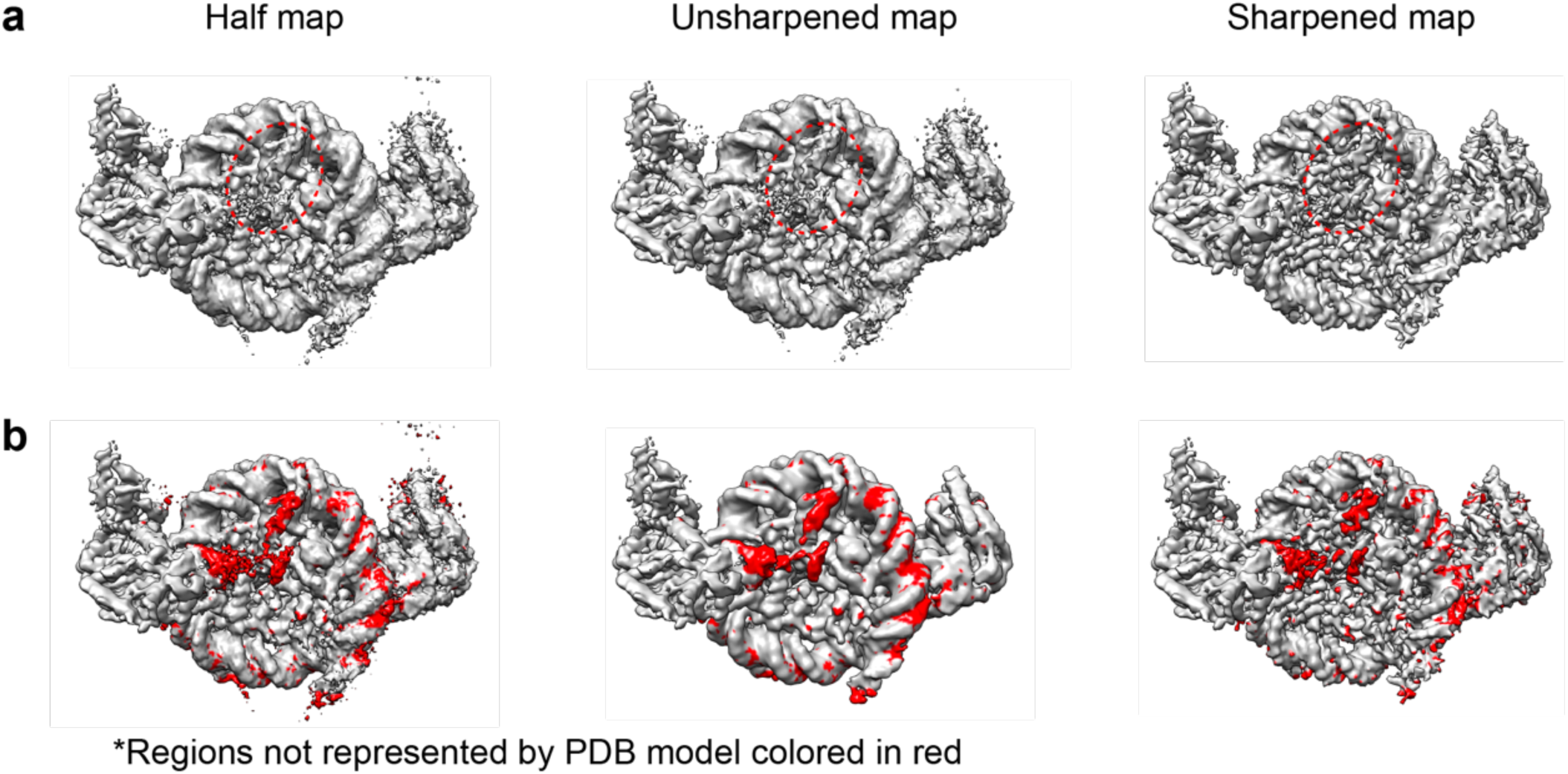
cryo-EM reconstructio of RSC^SAR^-nucleosome (ADP-BeF_3_) shows a region of unassigned density on the surface of the nucleosome. **a**, Different maps from the RSC^SAR^ consensus refinement are shown in the same orientation. A region of unassigned density is circled in red. **b**, The same maps shown in panel (a) with red indicating regions of the maps that are not accounted for by the PDB model of the nucleosome. For each map, a region of unassigned density is shown on the surface of the nucleosome.

**Supp.Fig.9.**
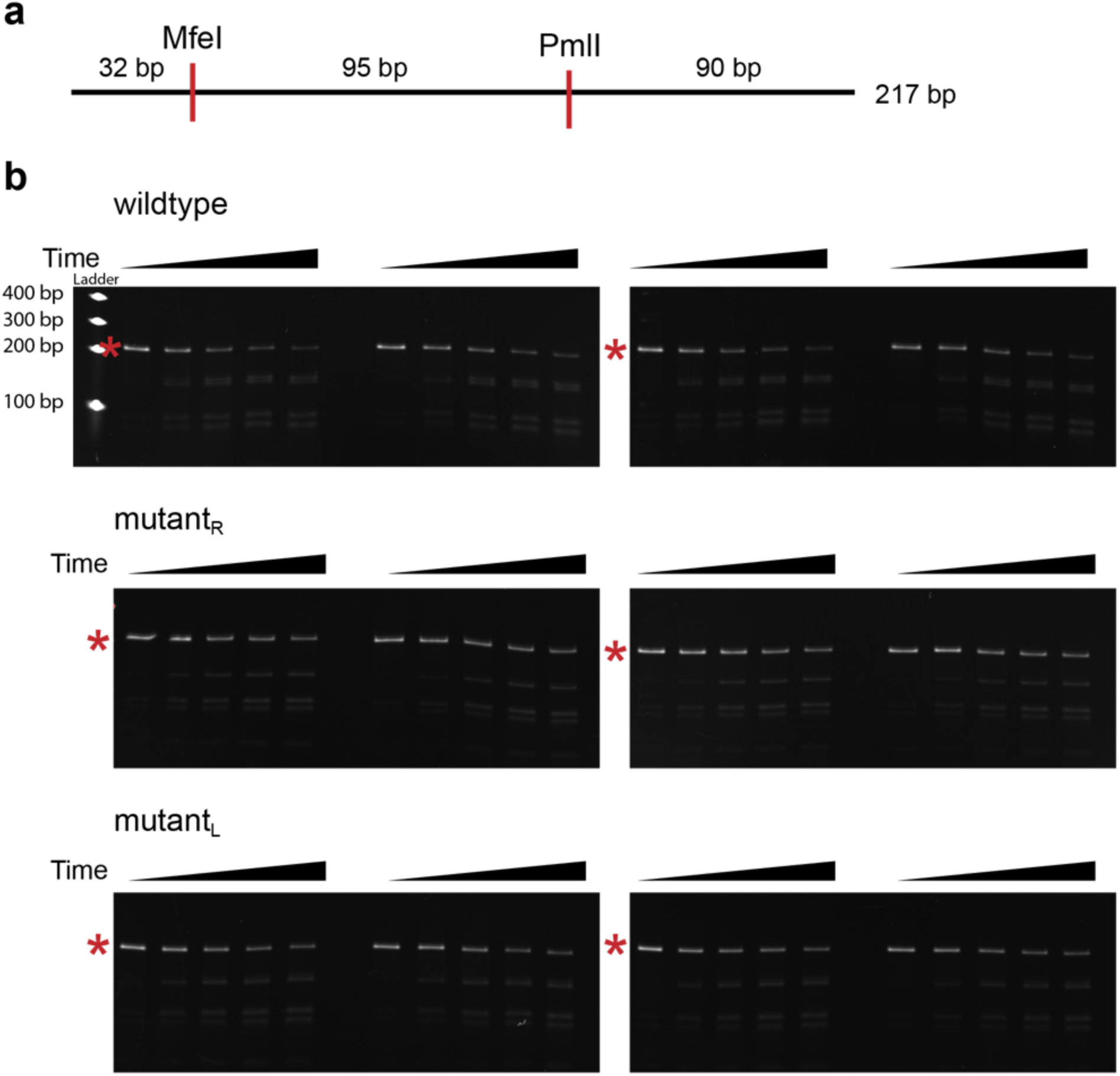
Remodeling assays show that a basic region in Sth1 enhances remodeling. **a**, DNA substrate used for the restriction enzyme accessibility assays: A 217 bp DNA fragment with the MfeI and PmlI restriction sites used in the assay highlighted. **b**, Remodeling assays were started with the addition of ATP. Aliquots were removed at 0, 5, 15, 30 and 45 minutes and incubated with Proteinase K to stop the reaction. Deproteinated samples were separated on a 10% TGX gel, stained with SYBR gold, and the uncleaved DNA band (*) was quantified by densitometry.

**Supplementary Table 1.**

|  | RSC<sup>SAR</sup><br>apo | RSC<sup>SAR</sup><br>nucleosome (BeF<sub>3</sub>) | RSC<sup>SAR</sup><br>nucleosome (BeF<sub>3</sub>)<br>Peeled DNA<br>conformation |
| --- | --- | --- | --- |
|  | PDB 6VZG<br>EMD-21489 | PDB 6VZ4<br>EMD-21484 | EMD-21493 |
| **Data Collection** |  |  |  |
| Microscope | Talos Arctica | Talos Arctica | Talos Arctica |
| Camera | K2 summit | K2 Summit | K2 Summit |
| Camera Mode | Super-Res | Counting | Counting |
| Magnification | 36,000 | 36,000 | 36,000 |
| Voltage (kV) | 200 | 200 | 200 |
| Phase Plate | Volta Phase Plate | none | none |
| Total electron exposure (e-/Å<sup>2</sup>) | 50-55 | 50-55 | 50-55 |
| Exposure rate (e-/pixel/sec) | 5-7.5 | 5-7.5 | 5-7.5 |
| Defocus Range (um) | 0.6-2.5 | 0.6-2.5 | 0.6-2.5 |
| Pixel Size (Å/pixel) | 0.58 | 1.16 | 0.58 |
| Micrographs collected (no.) | 5365 | 9292 | 9292 |
| **Reconstruction** |  |  |  |
| 3D Processing Package | Relion 3 | Relion 3 | Relion 3 |
| Total Extracted picks (no.) | 1,986,341 | 2,020,734 | 2,020,734 |
| Final Particles (no.) | 415,957 | 293,940 | 112,364 |
| Symmetry | C1 | C1 | C1 |
| Resolution (global) (Å) |  |  |  |
| FSC 0.143 (unmasked/masked) | (4.6/ 4.2) | (4.4/ 3.9) | (4.6/ 4.3) |
| Local resolution range (Å) | 4.0-5.8 | 3.5-15 |  |
| Map sharpening *B*-factor | -162 | -118 | -114 |
| **Refinement** |  |  |  |
| Model refinement package | Rosetta | Rosetta, Phenix | N/A |
| Number of models | 10 | 10 | N/A |
| Nonhydrogen atoms | 7420 (per model) | 23998 (per model) | N/A |
| Protein residues | 906 (per model) | 2511 (per model) | N/A |
| *B* factors (Å<sup>2</sup>) |  |  |  |
| Protein residues | 101.4 | 117.89 | N/A |
| R.m.s. deviations |  |  |  |
| Bond Lengths (Å) | 0.014 | 0.013 | N/A |
| Bond angles (°) | 1.35 | 1.12 | N/A |
| **Validation** |  |  |  |
| MolProbity score | 1.23 | 1.27 | N/A |
| Clashscore | 1.89 | 2.08 | N/A |
| Poor rotamers (%) | 0.01 | 0.04 | N/A |
| CaBLAM outliers (%) | 0 | 0 | N/A |
| Ramachandran plot |  |  |  |
| Favored (%) | 95.91 | 95.85 | N/A |
| Allowed (%) | 3.18 | 3.21 | N/A |
| Disallowed (%) | 0.91 | 0.94 | N/A |
| EMRinger score | 1.37 | 1.77 |  |
| Map CC (*CCmask*) | 0.791 | 0.832 | N/A |

## References

1. Bartholomew, B. Regulating the chromatin landscape: structural and mechanistic perspectives. Annual review of biochemistry 83, 671–696 (2014).

2. Clapier, C. R. & Cairns, B. R. The Biology of Chromatin Remodeling Complexes. Annual Review of Biochemistry 78, 273–304 (2009).

3. Kwon, H., Imbalzano, A. N., Khavari, P. A., Kingston, R. E. & Green, M. R. Nucleosome disruption and enhancement of activator binding by a human SW1/SNF complex. Nature 370, 477–481 (1994).

4. Parnell, T. J., Huff, J. T. & Cairns, B. R. RSC regulates nucleosome positioning at Pol II genes and density at Pol III genes. The EMBO journal 27, 100–110 (2008).

5. Cairns, B. R., Levinson, R. S., Yamamoto, K. R. & Kornberg, R. D. Essential role of Swp73p in the function of yeast Swi/Snf complex. Genes & Development 10, 2131–2144 (1996).

6. Stern, M., Jensen, R. & Herskowitz, I. Five SWI genes are required for expression of the HO gene in yeast. Journal of Molecular Biology 178, 853–868 (1984).

7. Kadoch, C. & Crabtree, G. R. Mammalian SWI/SNF chromatin remodeling complexes and cancer: Mechanistic insights gained from human genomics. Science advances 1, e1500447–e1500447 (2015).

8. Reisman, D., Glaros, S. & Thompson, E. A. The SWI/SNF complex and cancer. Oncogene 28, 1653–1668 (2009).

9. Shain, A. H. & Pollack, J. R. The spectrum of SWI/SNF mutations, ubiquitous in human cancers. PloS one 8, e55119–e55119 (2013).

10. Kadoch, C. et al. Proteomic and bioinformatic analysis of mammalian SWI/SNF complexes identifies extensive roles in human malignancy. Nature genetics 45, 592–601 (2013).

11. Wang, X. et al. Oncogenesis Caused by Loss of the SNF5 Tumor Suppressor Is Dependent on Activity of BRG1, the ATPase of the SWI/SNF Chromatin Remodeling Complex. Cancer Res 69, 8094 (2009).

12. Peterson, C. L., Zhao, Y. & Chait, B. T. Subunits of the Yeast SWI/SNF Complex Are Members of the Actin-related Protein (ARP) Family. J. Biol. Chem. 273, 23641–23644 (1998).

13. Zhao, K. et al. Rapid and Phosphoinositol-Dependent Binding of the SWI/SNF-like BAF Complex to Chromatin after T Lymphocyte Receptor Signaling. Cell 95, 625–636 (1998).

14. Cairns, B. R., Erdjument-Bromage, H., Tempst, P., Winston, F. & Kornberg, R. D. Two Actin-Related Proteins Are Shared Functional Components of the Chromatin-Remodeling Complexes RSC and SWI/SNF. Molecular Cell 2, 639–651 (1998).

15. Szerlong, H. et al. The HSA domain binds nuclear actin-related proteins to regulate chromatin-remodeling ATPases. Nature structural & molecular biology 15, 469–476 (2008).

16. Schubert, H. L. et al. Structure of an actin-related subcomplex of the SWI/SNF chromatin remodeler. Proceedings of the National Academy of Sciences 110, 3345–3350 (2013).

17. Turegun, B., Baker, R. W., Leschziner, A. E. & Dominguez, R. Actin-related proteins regulate the RSC chromatin remodeler by weakening intramolecular interactions of the Sth1 ATPase. Communications biology 1, 1–1 (2018).

18. Clapier, C. R. et al. Regulation of DNA Translocation Efficiency within the Chromatin Remodeler RSC/Sth1 Potentiates Nucleosome Sliding and Ejection. Molecular cell 62, 453–461 (2016).

19. Liu, X., Li, M., Xia, X., Li, X. & Chen, Z. Mechanism of chromatin remodelling revealed by the Snf2-nucleosome structure. Nature 544, 440 (2017).

20. Patel, A. B. et al. Architecture of the chromatin remodeler RSC and insights into its nucleosome engagement. eLife 8, e54449 (2019).

21. Wagner, F. R. et al. Structure of SWI/SNF chromatin remodeller RSC bound to a nucleosome. Nature (2020) doi: 10.1038/s41586-020-2088-0.

22. Ye, Y. et al. Structure of the RSC complex bound to the nucleosome. Science eaay0033 (2019) doi: 10.1126/science.aay0033.

23. Han, Y., Reyes, A. A., Malik, S. & He, Y. Cryo-EM structure of SWI/SNF complex bound to a nucleosome. Nature (2020) doi: 10.1038/s41586-020-2087-1.

24. He, S. et al. Structure of nucleosome-bound human BAF complex. Science 367, 875 (2020).

25. Eastlund, A., Malik, S. S. & Fischer, C. J. Kinetic mechanism of DNA translocation by the RSC molecular motor. Archives of Biochemistry and Biophysics 532, 73–83 (2013).

26. Malik, S. S., Rich, E., Viswanathan, R., Cairns, B. R. & Fischer, C. J. Allosteric Interactions of DNA and Nucleotides with S. cerevisiae RSC. Biochemistry 50, 7881–7890 (2011).

27. Sirinakis, G. et al. The RSC chromatin remodelling ATPase translocates DNA with high force and small step size. The EMBO Journal 30, 2364–2372 (2011).

28. Sirinakis, G. et al. The RSC chromatin remodelling ATPase translocates DNA with high force and small step size. The EMBO Journal 30, 2364–2372 (2011).

29. Li, M. et al. Mechanism of DNA translocation underlying chromatin remodelling by Snf2. Nature 567, 409–413 (2019).

30. Xia, X., Liu, X., Li, T., Fang, X. & Chen, Z. Structure of chromatin remodeler Swi2/Snf2 in the resting state. Nature Structural and Molecular Biology 23, nsmb.3259 (2016).

31. Nakane, T., Kimanius, D., Lindahl, E., Scheres, S. H. & Brunger, A. T. Characterisation of molecular motions in cryo-EM single-particle data by multi-body refinement in RELION. eLife 7, e36861 (2018).

32. Armache, K.-J., Garlick, J. D., Canzio, D., Narlikar, G. J. & Kingston, R. E. Structural Basis of Silencing: Sir3 BAH Domain in Complex with a Nucleosome at 3.0 A Resolution. Science 334, 977–982 (2011).

33. Barbera, A. J. et al. The Nucleosomal Surface as a Docking Station for Kaposi’s Sarcoma Herpesvirus LANA. Science 311, 856 (2006).

34. Kato, H. et al. A Conserved Mechanism for Centromeric Nucleosome Recognition by Centromere Protein CENP-C. Science 340, 1110–1113 (2013).

35. Makde, R. D., England, J. R., Yennawar, H. P. & Tan, S. Structure of RCC1 chromatin factor bound to the nucleosome core particle. Nature 467, 562–566 (2010).

36. Mashtalir, N. et al. Modular Organization and Assembly of SWI/SNF Family Chromatin Remodeling Complexes. Cell 175, 1272–1288.e20 (2018).

37. Levendosky, R. F. & Bowman, G. D. Asymmetry between the two acidic patches dictates the direction of nucleosome sliding by the ISWI chromatin remodeler. eLife 8, e45472 (2019).

38. Dann, G. P. et al. ISWI chromatin remodellers sense nucleosome modifications to determine substrate preference. Nature 548, 607 (2017).

39. Dao, H. T., Dul, B. E., Dann, G. P., Liszczak, G. P. & Muir, T. W. A basic motif anchoring ISWI to nucleosome acidic patch regulates nucleosome spacing. Nature Chemical Biology 16, 134–142 (2020).

40. Valencia, A. M. et al. Recurrent SMARCB1 Mutations Reveal a Nucleosome Acidic Patch Interaction Site That Potentiates mSWI/SNF Complex Chromatin Remodeling. Cell 179, 1342–1356.e23 (2019).

41. Turegun, B., Baker, R. W., Leschziner, A. E. & Dominguez, R. Actin-related proteins regulate the RSC chromatin remodeler by weakening intramolecular interactions of the Sth1 ATPase. Communications biology 1, 1–1 (2018).

42. Turegun, B., Kast, D. J. & Dominguez, R. Subunit Rtt102 controls the conformation of the Arp7/9 heterodimer and its interactions with nucleotide and the catalytic subunit of SWI/SNF remodelers. The Journal of biological chemistry 288, 35758–35768 (2013).

43. Dann, G. P. et al. ISWI chromatin remodellers sense nucleosome modifications to determine substrate preference. Nature 548, 607 (2017).

44. Dyer, P. N. et al. Reconstitution of Nucleosome Core Particles from Recombinant Histones and DNA. in vol. 375 23–44 (2003).

45. Stark, H. Chapter Five GraFix: Stabilization of Fragile Macromolecular Complexes for Single Particle Cryo-EM. Methods in Enzymology 481, 109–126 (2010).

46. Herzik, J. M. A., Wu, M. & Lander, G. C. Achieving better-than-3-Å resolution by single-particle cryo-EM at 200 keV. Nature methods 14, 1075–1078 (2017).

47. Lander, G. C. et al. Appion: an integrated, database-driven pipeline to facilitate EM image processing. Journal of structural biology 166, 95–102 (2009).

48. Kremer, J. R., Mastronarde, D. N. & McIntosh, J. R. Computer Visualization of Three-Dimensional Image Data Using IMOD. Journal of Structural Biology 116, 71–76 (1996).

49. Zheng, S. Q. et al. MotionCor2: anisotropic correction of beam-induced motion for improved cryo-electron microscopy. Nature Methods 14, 331–332 (2017).

50. Rohou, A. & Grigorieff, N. CTFFIND4: Fast and accurate defocus estimation from electron micrographs. Journal of Structural Biology 192, 216–221 (2015).

51. Zivanov, J. et al. New tools for automated high-resolution cryo-EM structure determination in RELION-3. eLife 7, e42166 (2018).

52. Wagner, T. et al. SPHIRE-crYOLO is a fast and accurate fully automated particle picker for cryo-EM. Communications Biology 2, 218 (2019).

53. Punjani, A., Rubinstein, J. L., Fleet, D. J. & Brubaker, M. A. cryoSPARC: algorithms for rapid unsupervised cryo-EM structure determination. Nature Methods 14, 290–296 (2017).

54. Schubert, H. L. et al. Structure of an actin-related subcomplex of the SWI/SNF chromatin remodeler. Proceedings of the National Academy of Sciences 110, 3345–3350 (2013).

55. Emsley, P. & Cowtan, K. Coot: model-building tools for molecular graphics. Acta Crystallographica Section D: Biological Crystallography 60, 2126–2132 (2004).

56. Cianfrocco, M. A., Lahiri, I., DiMaio, F. & Leschziner, A. E. cryoem-cloud-tools: A software platform to deploy and manage cryo-EM jobs in the cloud. Journal of Structural Biology 203, 230–235 (2018).

57. Song, Y. et al. High-resolution comparative modeling with RosettaCM. Structure (London, England : 1993) 21, 1735–1742 (2013).

58. Chen, V. B. et al. MolProbity: all-atom structure validation for macromolecular crystallography. Acta crystallographica. Section D, Biological crystallography 66, 12–21 (2010).

59. Nakane, T., Kimanius, D., Lindahl, E. & Scheres, S. H. Characterisation of molecular motions in cryo-EM single-particle data by multi-body refinement in RELION. eLife 7, e36861 (2018).

60. Xia, X., Liu, X., Li, T., Fang, X. & Chen, Z. Structure of chromatin remodeler Swi2/Snf2 in the resting state. Nature Structural and Molecular Biology 23, nsmb.3259 (2016).

61. Alford, R. F. et al. The Rosetta All-Atom Energy Function for Macromolecular Modeling and Design. Journal of chemical theory and computation 13, 3031–3048 (2017).

62. Afonine, P. V. et al. Real-space refinement in PHENIX for cryo-EM and crystallography. Acta crystallographica. Section D, Structural biology 74, 531–544 (2018).

63. Liebschner, D. et al. Macromolecular structure determination using X-rays, neutrons and electrons: recent developments in Phenix. Acta Crystallographica Section D 75, 861—877 (2019).

